# Hybrid misexpression in multiple developing tissues within a recent adaptive radiation of *Cyprinodon* pupfishes

**DOI:** 10.1101/372912

**Authors:** Joseph A. McGirr, Christopher H. Martin

**Affiliations:** Department of Biology, University of North Carolina at Chapel Hill, 1250 South Rd., Chapel Hill, NC 27514-3280

**Keywords:** RNA-seq, F_1_ hybrid, misexpression, trophic specialization, allele specific expression, adaptive radiation, transcriptomics

## Abstract

Genetic incompatibilities constitute the final stages of reproductive isolation and speciation, but little is known about incompatibilities that occur within recent adaptive radiations among closely related diverging populations. Crossing divergent species to form hybrids can break up coadapted variation, resulting in genetic incompatibilities within developmental networks shaping adaptive traits. We crossed two closely related sympatric *Cyprinodon* pupfish species – a dietary generalist and a specialized molluscivore – and measured expression levels in their F_1_ hybrids to identify regulatory variation underlying the novel craniofacial morphology found in this recent microendemic adaptive radiation. We extracted mRNA from eight day old whole-larvae tissue and from craniofacial tissues dissected from 17-20 day old larvae to compare gene expression between a total of seven F_1_ hybrids and 24 individuals from parental species populations. We found 3.9% of genes differentially expressed between generalists and molluscivores in whole-larvae tissues and 0.6% of genes differentially expressed in craniofacial tissue. We found that 2.1% of genes were misexpressed in whole-larvae hybrids at 8 dpf whereas 19.1% of genes were misexpressed in hybrid craniofacial tissue at 17-20 dpf, after correcting for sequencing biases. We also measured allele specific expression across 15,429 phased heterozygous sites to identify regulatory mechanisms underlying differential expression between generalists and molluscivores. Together, our results highlight the importance of considering misexpression as an early indicator of genetic incompatibilities in the context of rapidly diverged morphology and suggests that wide-spread compensatory regulatory divergence drives hybrid misexpression in developing tissues that give rise to novel craniofacial traits.

## Introduction

Changes in gene expression are an important source of variation in adaptive morphological traits (Carroll 2008; Wolf et al. 2010; Indjeian et al. 2016). As genetic variation accumulates in regulatory and coding sequences, stabilizing selection on gene expression results in coevolution such that molecular functions are largely maintained (Coolon et al. 2014; Hodgins-Davis et al. 2015). Crossing divergent species to form F_1_ hybrids can break up such coadapted variation, resulting in genetic incompatibilities within developing tissues that give rise to adaptive traits (Michalak and Noor 2004; Landry et al. 2007; Mack and Nachman 2017). Genetic incompatibilities that reduce hybrid fitness can drive reproductive isolation either intrinsically – causing sterility or increased embryonic mortality – or extrinsically whereby incompatibilities reduce hybrid performance in a particular environment (Schluter 2000, Coyne and Orr 2004).

Of particular importance to the process of speciation are genetic incompatibilities caused by hybrid misexpression – when gene expression levels in hybrids are transgressive and fall outside of the range of expression variation observed in both parental species (Michalak and Noor 2004; Ranz et al. 2004; Haerty and Singh 2006; Rockman and Kruglyak 2006; Malone and Michalak 2008; Renaut et al. 2009). This pattern of expression causes Dobzhansky-Muller incompatibilities (DMIs) if incompatible alleles in hybrids cause misexpression that results in reduced hybrid fitness and thus increased post-zygotic reproductive isolation (Presgraves 2003; Coyne 2004; Sweigart et al. 2006; Ortíz-Barrientos et al. 2007; Malone and Michalak 2008; Renaut et al. 2009; Davidson and Balakrishnan 2016). Laboratory studies searching for genes that cause DMIs often identify genes causing sterility or embryonic lethality in hybrids. This approach ignores the fitness consequences of misexpression occurring at later developmental stages within diverse tissue types, thus underestimating the actual number of genetic incompatibilities distinguishing species (Fang et al. 2012; Schumer et al. 2014). Combining findings from these studies with analyses of hybrid misexpression in tissues that give rise to adaptive morphological traits can reveal a broader view of incompatibilities that arise during speciation.

Studies of gene expression in hybrids can also implicate regulatory mechanisms underlying expression divergence between parental species, which is important for understanding how expression levels are inherited and how they shape adaptive traits (Wittkopp et al. 2004; McManus et al. 2010; Mack and Nachman 2017). Research on hybrid gene expression thus far has shown mixed results regarding patterns of inheritance (Signor and Nuzhdin 2018). Some studies found evidence for ubiquitous transgressive expression inherited in F_1_ hybrids (i.e. over- or under-dominance) (Ranz et al. 2004; Rockman and Kruglyak 2006; Roberge et al. 2008), while others found predominately additive patterns (Hughes et al. 2006; Rottscheidt and Harr 2007; Davidson and Balakrishnan 2016). Mechanisms of gene expression divergence in F_1_ hybrids are characterized as interactions between *cis*-regulatory elements and *trans*-regulatory factors. *Cis* elements are often non-coding regions of DNA proximal to genes that are bound by *trans*-acting proteins and RNAs to regulate mRNA abundance. It is possible to identify mechanisms of gene expression divergence between parental species by bringing *cis* elements from both parents together in the same *trans* environment in F_1_ hybrids and quantifying allele specific expression (ASE) of parental alleles at heterozygous sites (Cowles et al. 2002; Wittkopp et al. 2004). *Cis* and *trans* regulatory variants can compensate for one another if stabilizing selection favors an optimal level of gene expression. Hybrid misexpression is expected when different compensatory variants have accumulated in diverging lineages (Denver et al. 2005; Landry et al. 2005; Bedford and Hartl 2009; Goncalves et al. 2012).

Here we investigate F_1_ hybrids from crosses between two closely related species of *Cyprinodon* pupfishes to understand regulatory mechanisms that led to the evolution of novel craniofacial adaptations in this group (Fig 1A). *Cyprinodon brontotheroides* – hereafter referred to as the molluscivore – is a trophic specialist species endemic to San Salvador Island, Bahamas that has adapted to eat hard shelled prey including mollusks and ostracods (Martin and Wainwright 2013a,c). This species likely diverged from a generalist common ancestor within the past 10,000 years to occupy this novel niche (Mylroie, J.E, Hagey 1995; Holtmeier 2001; Turner et al. 2008; Martin and Wainwright 2011; Martin 2016). Adapting to this niche involved extreme morphological divergence in craniofacial traits compared to its sympatric generalist sister species *Cyprinodon variegatus* (Martin and Wainwright 2013c; Lencer et al. 2016). This species consumes mainly algae and detritus and is hereafter referred to as the ‘generalist.’ Almost all other Caribbean pupfish species are generalists, with the exception of a novel scale-eating pupfish that is also a member of the San Salvador pupfish radiation (Martin and Wainwright 2011, 2013c) and a second sympatric radiation of trophic specialists in Laguna Chichancanab, Mexico (Humphries and Miller 1981; Strecker 2006). Molluscivores exhibit a novel skeletal protrusion on the anteriodorsal head of the maxilla not found in generalist populations that may be used to stabilize prey items held within its oral jaws, which are shorter and more robust relative to generalist species (Fig 1A). This jaw morphology provides higher mechanical advantage for crushing mollusks and other hard-shelled prey (Wainwright and Richard 1995; Martin and Wainwright 2011).

**Fig 1.**
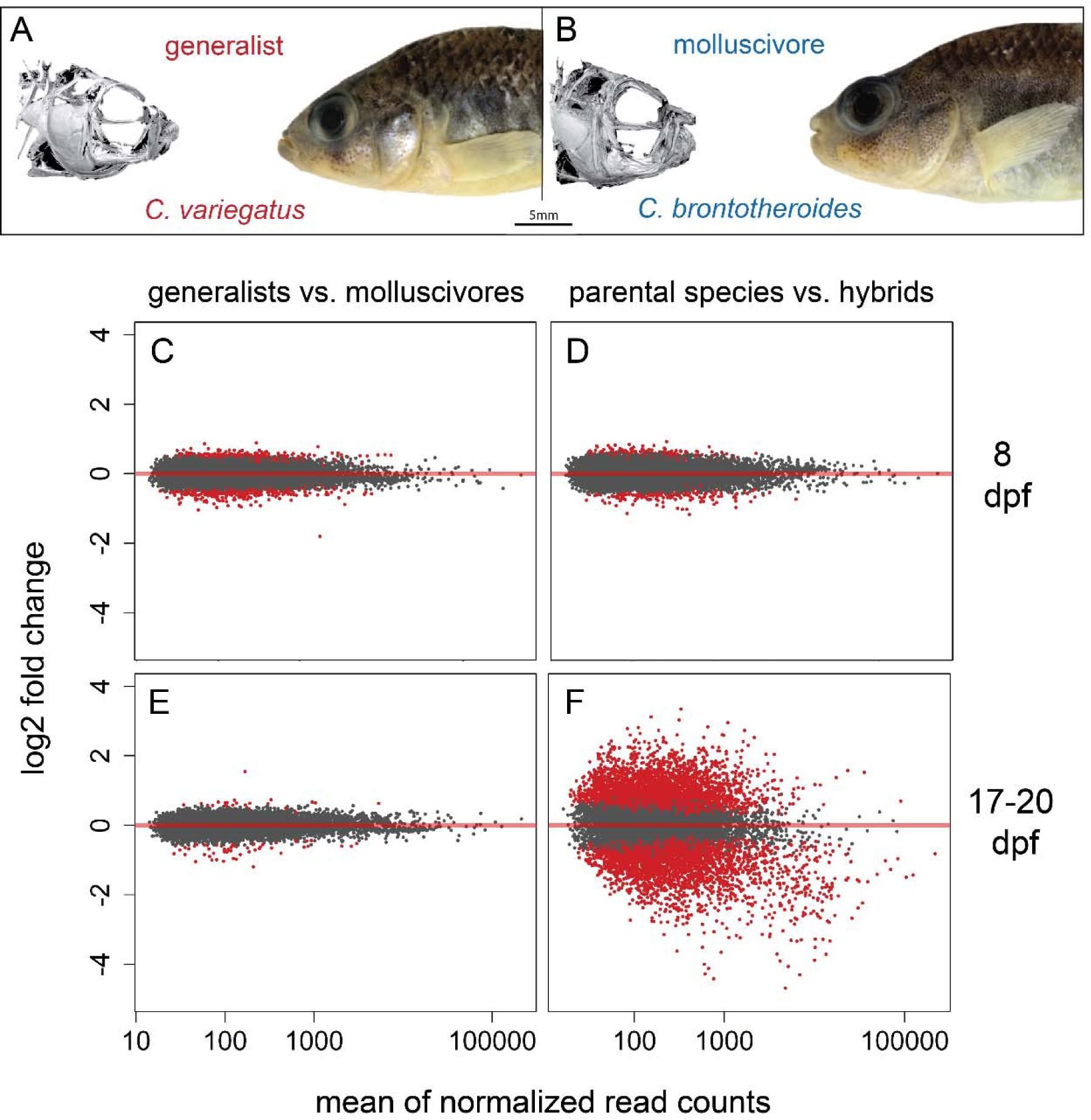
Extensive misexpression in F_1_ hybrid craniofacial tissues. A) *Cyprinodon variegatus* – the generalist. B) *C. brontotheroides* – the molluscivore (µCT scans of the cranial skeleton of each species modified from Hernandez et al. 2018). Variation in gene expression between generalists vs. molluscivores at 8 dpf, D) parental species vs. hybrids at 8 dpf, E) generalists vs. molluscivores at 17-20 dpf, and F) parental species vs. hybrids at 17-20 dpf. Red points indicate genes detected as differentially expressed at 5% false discovery rate with Benjamini-Hochberg multiple testing adjustment. Grey points indicate genes showing no significant difference in expression between groups. Red line indicates a log_2_ fold change of zero between groups. Points above/below the line are upregulated/downregulated in molluscivores relative to generalists (C and E) or hybrids relative to parental species (D and F).

Molluscivores and generalists readily hybridize in the laboratory to produce fertile F_1_ offspring with intermediate craniofacial morphologies and no obvious sex ratio distortion (Martin and Wainwright 2011, 2013b; Martin and Feinstein 2014). These species remain largely reproductively isolated in sympatry across multiple lake populations (genome-wide average F_st_ = 0.08; (Martin and Feinstein 2014; West and Kodric-Brown 2015; McGirr and Martin 2017). Therefore, unlike most studies of hybrid misexpression, we are not solely concerned with identifying gene expression patterns underlying hybrid sterility or lethality. Rather, we also aim to characterize misexpression in a developing tissue that gives rise to novel craniofacial phenotypes within a young species pair with ongoing gene flow. We dissected craniofacial tissue from 17-20 day old F_1_ hybrids and extracted total mRNA to quantify gene expression levels. We also extracted whole-larvae mRNA from 8 day old generalists, molluscivores, and their hybrids. We found misexpression in hybrids at both stages. Finally, we quantified allele specific expression (ASE) across exome-wide phased heterozygous sites to uncover mechanisms of regulatory divergence and found evidence for compensatory variation influencing patterns of hybrid misexpression.

## Materials and Methods

### Study system and sample collection

Our methods for raising larvae and extracting RNA were identical to previously outlined methods (McGirr and Martin 2018). We collected fishes for breeding from three hypersaline lakes on San Salvador Island, Bahamas (Little Lake, Osprey Lake, and Crescent Pond) using a hand net or seine net between 2011 and 2017. These fishes were reared at 25–27°C, 10–15 ppt salinity, pH 8.3, and fed a mix of commercial pellet foods and frozen foods. All lab bred larvae were raised exclusively on newly hatched brine shrimp after hatching and before euthanasia.

Individuals were euthanized in an overdose of buffered MS 222 and stored in RNA later (Ambion, Inc.) at −20°C for up to 18 months. We used RNeasy Mini Kits (Qiagen catalog #74104) to extract RNA from all samples.

We previously generated 24 transcriptomes belonging to generalists and molluscivores collected at two early developmental stages: 8-10 days post fertilization and 17-20 dpf (McGirr and Martin 2018). RNA was extracted from whole-larvae tissue at 8-10 dpf. We dissected all 17-20 dpf samples to extract RNA from anterior craniofacial tissues containing the dentary, angular, articular, maxilla, premaxilla, palatine, and associated craniofacial connective tissues (Fig. S1). Dissections were performed using fine tipped tweezers washed with RNase AWAY (Molecular BioProducts). These 24 samples were generated by breeding populations of lab-raised fishes that resulted from either one or two generations of full-sib breeding between wild caught fishes from Little Lake and Crescent Pond on San Salvador Island, Bahamas (Table 1). There was variation in sampling time because eggs were fertilized naturally within breeding tanks and collected on the same day or subsequent day following egg laying. We collected larvae in a haphazard manner over multiple spawning intervals and it is unlikely that sampling time varied consistently by species.

**Table 1.** Sampling design for mRNA sequencing. Parental fishes crossed to produce larvae for sequencing were either wild-caught (F_0_) or lab-raised over *n* generations (indicated by F*_n_*).

| mothers | fathers | offspring<br>sampled | lake<br>population | sequencing<br>round | stage<br>(dpf) |
| --- | --- | --- | --- | --- | --- |
| $F_0$ generalist | $F_0$ molluscivore | 3 hybrids | osprey lake | 4 | 8 |
| $F_0$ generalists | $F_0$ generalists | 3 generalists | osprey lake | 3 | 8 |
| $F_0$ molluscivores | $F_0$ molluscivores | 3 molluscivores | osprey lake | 3 | 8 |
| $F_0$ generalists | $F_0$ generalists | 3 generalists | crescent pond | 3 | 8 |
| $F_0$ molluscivores | $F_0$ molluscivores | 3 molluscivore | crescent pond | 4 | 8 |
| $F_1$ generalists | $F_1$ generalists | 3 generalists | little lake | 1 | 8-10 |
| $F_2$ molluscivores | $F_2$ molluscivores | 3 molluscivores | little lake | 1 | 8-10 |
| $F_2$ generalists | $F_2$ generalists | 3 generalists | crescent pond | 1 | 8-10 |
| $F_2$ molluscivores | $F_2$ molluscivores | 3 molluscivores | crescent pond | 1 | 8-10 |
| $F_2$ generalist | $F_3$ molluscivore | 4 hybrids | little lake | 2 | 17-20 |
| $F_1$ generalists | $F_1$ generalists | 3 generalists | little lake | 1 | 17-20 |
| $F_2$ molluscivores | $F_2$ molluscivores | 3 molluscivores | little lake | 1 | 17-20 |
| $F_2$ generalists | $F_2$ generalists | 3 generalists | crescent pond | 1 | 17-20 |
| $F_2$ molluscivores | $F_2$ molluscivores | 3 molluscivores | crescent pond | 1 | 17-20 |

Here we analyze an additional 19 transcriptomes from generalists, molluscivores, and their hybrids (Table 1). First, we crossed a generalist female with a molluscivore male to generate four F_1_ hybrids that were collected at 17-20 dpf and extracted RNA from dissected craniofacial tissues. A lab-reared female generalist was used to generate hybrids that was derived from wild caught generalists from Little Lake following one generation of full-sib mating. A lab-reared male molluscivore was used to generate hybrids that was derived from wild caught molluscivores from Little Lake following two generations of full-sib mating.

We performed separate crosses to collect larvae at exactly 8 dpf (190-194 hours after fertilization rather than 8-10 days). We crossed a generalist female with a molluscivore male to generate three F_1_ hybrids for whole-larvae RNA extractions. The parents of these hybrids were wild-caught from Osprey Lake. Finally, we extracted whole-larvae RNA from six generalists and six molluscivores collected at 8 dpf. These samples were generated from wild-caught individuals from Osprey Lake and Crescent Pond. In total, we analyzed transcriptomes from 43 individuals that involved four separate rounds of sequencing (Table 1 and S1).

### RNA sequencing and alignment

The previously reported 24 transcriptomes were sequenced at the High Throughput Genomic Sequencing Facility at UNC Chapel Hill in April 2017 (McGirr and Martin 2018). The 24 libraries were prepared at the facility using the KAPA stranded mRNA seq kit (KAPA Biosystems 2016) followed by sequencing on one lane of Illumina 150 paired-end Hiseq4000 (Table 1 and 2).

19 additional transcriptomes were sequenced at The Vincent J. Coates Genomics Sequencing Laboratory at the University of California, Berkeley. All 19 libraries were prepared at the facility using the Illumina stranded Truseq RNA kit (Illumina RS-122-2001) and all sequencing was performed on Illumina 150 paired-end Hiseq4000. Four libraries for RNA extracted from 17-20 dpf hybrid craniofacial tissues were pooled on a single lane and sequenced in June 2017. 15 libraries for whole-larvae RNA samples collected at exactly 8 dpf were pooled across one and three lanes and sequenced in May (n = 9) and July (n = 6) 2018, respectively (Table 1 and S1).

We filtered all raw reads using Trim Galore (v. 4.4, Babraham Bioinformatics) to remove Illumina adaptors and low quality reads (mean Phred score < 20) and mapped filtered reads to the scaffolds of the *Cyprinodon* reference genome (NCBI, *C. variegatus* annotation release 100, total sequence length = 1,035,184,475; number of scaffolds = 9259, scaffold N50 = 835,301; contig N50 = 20,803; (Lencer et al. 2017)) using the RNA seq aligner STAR with default parameters (v. 2.5 (Dobin et al. 2013)). We used the featureCounts function of the Rsubread package (Liao et al. 2014) requiring paired end and reverse stranded options to generate read counts across 24,952 previously annotated features (Lencer et al. 2017) with an average coverage depth of 136 reads (Table S2 and S3). We assessed mapping and count quality using MultiQC (Ewels et al. 2016). We previously showed that there was no difference between generalists and molluscivores in the proportion of reads that map to annotated features of the *Cyprinodon* reference genome (McGirr and Martin 2018). Similarly, here we found no difference in the proportion of reads mapping to features between generalists, molluscivores, and hybrids (Fig. S2; ANOVA, *P* = 0.6), but we did find that fewer reads mapped to features in 17-20 dpf samples than 8 dpf samples (ANOVA, *P* = 2.38 × 10^−10^).

Since we analyzed RNA from 43 individuals that were sequenced across four different dates and their libraries were prepared with either KAPA or TruSeq stranded mRNA-seq kits, we tested whether a significant amount of between-sample variance in read counts was explained by sequencing date or library preparation kit. We fit linear models (using the lm() function in R) to determine whether normalized counts across individuals were influenced by 1) the number of PCR duplicate reads produced during sequence amplification, 2) the uniformity of coverage across a transcript, or 3) the depth of coverage across GC-rich transcripts. All of these measures could have been influenced by different library preparation methods (Alberti et al. 2014; Van Dijk et al. 2014; KAPA Biosystems 2016). We quantified the number of duplicate reads and the median percent GC content of mapped reads for each sample using RSeQC (Wang et al. 2012). We also used RSeQC to estimate transcript integrity numbers (TINs) which is a measure of potential *in vitro* RNA degradation within a sample. TIN is calculated by analyzing the uniformity of coverage across transcripts. (Wang et al. 2012, 2016). We performed ANOVA to determine whether the proportion of duplicate reads, GC content of reads, TINs, the number of normalized read counts, number of raw read counts, or number of raw fastq reads differed between samples grouped by library preparation method and by sequencing date.

### Differential expression analyses and hybrid inheritance of expression patterns

We performed differential expression analyses with DESeq2 (v. 3.5 (Love et al. 2014)). This program fits negative binomial generalized linear models for each gene across samples to test the null hypothesis that the fold change in gene expression between two groups is zero. DESeq2 uses an empirical Bayes shrinkage method to determine gene dispersion parameters, which models within-group variability in gene expression, and logarithmic fold changes in gene expression. DESeq2 normalizes raw read counts by calculating a geometric mean of counts for each gene across samples, dividing individual gene counts by this mean, and using the median of these ratios as a size factor for each sample. These sample-specific size factors account for differences in library size and sequencing depth between samples. Gene features showing less than 10 normalized counts in every sample in each comparison were discarded from analyses. Differential expression between groups was determined with Wald tests by comparing normalized posterior log fold change estimates and correcting for multiple testing using the Benjamini–Hochberg procedure with a false discovery rate of 0.05 (Benjamini and Hochberg 1995). We also used DESeq2 to perform clustering and principal component analyses (Fig. S3).

We conducted pairwise comparisons to identify genes differentially expressed between hybrids vs. parental species, hybrids vs. generalists, hybrids vs. molluscivores, and generalists vs. molluscivores. “Parental species” refers to generalists and molluscivores derived from the same populations as the parents of the hybrid samples. We did not sequence any of the parents crossed to generate hybrids. We defined genes as misexpressed in hybrids if they were significantly differentially expressed between hybrids and the parental species samples. First, we compared whole-larvae gene expression between samples collected at 8 dpf (six generalists, six molluscivores, and three hybrids). All of the 8 dpf samples were sequenced at the Vincent J. Coates Genomic Sequencing Laboratory, University of California Berkeley (VJCGSL UCB) and their libraries were all prepared using the TruSeq stranded mRNA-seq kit. Second, we compared craniofacial tissue gene expression between samples collected at 17-20 dpf (six generalists, six molluscivores, and four hybrids). The generalist and molluscivore samples were sequenced at the High-Throughout Sequencing Facility, University of North Carolina Chapel Hill (HTSF UNC) and their libraries were prepared using the KAPA stranded mRNA-seq kit, while the hybrids were sequenced at the VJCGSL UCB and their libraries were prepared using the TruSeq kit. In order to understand how sequencing at different facilities and using different library prep methods affected the proportion of genes misexpressed between hybrids and parental species at 17-20 dpf, we performed a third set of comparisons between hybrids collected at 8 dpf (sequenced at VJCGSL UCB with TruSeq) and generalists and molluscivores from a previous study collected at 8-10 dpf (sequenced at HTSF UNC with KAPA; (McGirr and Martin 2018)). We measured how many genes were differentially expressed between 8 dpf hybrids vs. 8-10 dpf parental species than there were differentially expressed between 8 dpf hybrids vs. 8 dpf parental species. This allowed us to estimate an upper-limit on the proportion of genes falsely identified as differentially expressed between 17-20 dpf hybrids and 17-20 dpf parental species due to samples being sequenced at different facilities with different library preparation kits.

To determine whether genes showed additive, dominant, or transgressive patterns of inheritance, we quantified differences in gene expression between hybrids vs. parental species and compared them to genes differentially expressed between generalists vs. molluscivores (Fig. 2). Hybrid inheritance was considered additive if hybrid gene expression was intermediate between generalists and molluscivores with significant differential expression between generalists and molluscivores, respectively. Inheritance was dominant if hybrid expression was significantly different from one parent species but not the other. Genes showing misexpression in hybrids showed transgressive inheritance, meaning hybrid gene expression was significantly higher (overdominant) or lower (underdominant) than both parental species.

**Fig 2.**
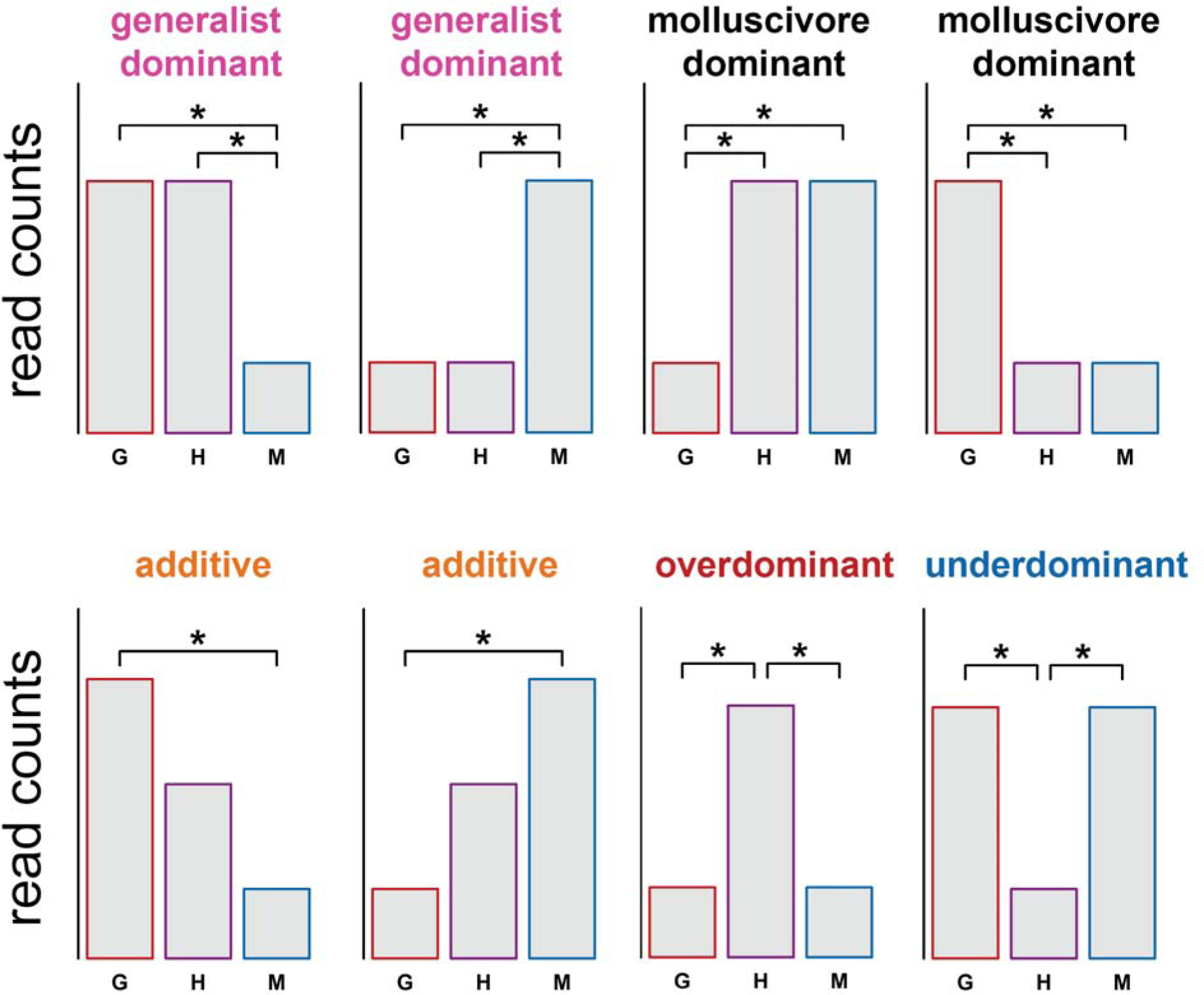
Classifying gene expression inheritance in hybrids. Schematic showing how gene expression inheritance in hybrids was classified. Asterisks indicate significant differential expression between groups. G = generalists, H = hybrids, M = molluscivores.

### Gene ontology enrichment analyses

The *Cyprinodon* reference genome is annotated for genomic features (NCBI, *C. variegatus* Annotation Release 100, (Lencer et al. 2017)), and many annotated genes share the same name as their zebrafish orthologs. We performed gene ontology (GO) enrichment analyses for genes differentially expressed between species and misexpressed in hybrids that shared the same name as zebrafish orthologs using GO Consortium resources available at geneontology.org (Ashburner et al. 2000; GO Consortium 2017). We searched for enrichment across biological process ontologies curated for zebrafish.

### Allele specific expression and mechanisms of regulatory divergence

We followed the best practices guide recommended by the Genome Analysis Toolkit (DePristo et al. 2011 (v. 3.5)) in order to call and refine SNP variants within coding gene regions using the Haplotype Caller function. We called SNPs across all filtered reads mapped to annotated features for 17-20 dpf samples and 8 dpf samples using conservative hard-filtering parameters (DePristo et al. 2011; Marsden et al. 2014): Phred-scaled variant confidence divided by the depth of nonreference samples > 2.0, Phred-scaled *P*-value using Fisher’s exact test to detect strand bias > 60, Mann–Whitney rank-sum test for mapping qualities (z > 12.5), Mann–Whitney rank-sum test for distance from the end of a read for those with the alternate allele (z > 8.0). We used the ReadBackedPhasing function with a phase quality threshold > 20 to identify phased variants. We used the VariantsToTable function (with genotypeFilterExpression “isHet = = 1”) to output phased heterozygous variants for each individual. We counted the number of reads covering heterozygous sites using the ASEReadCounter (with –U ALLOW_N_CIGAR_READS-minDepth 20 –minMappingQuality 10 –minBaseQuality 20-drf DuplicateRead). In total we identified 15,429 phased heterozygous sites across all 32 individuals with sequencing coverage ≥ 20× that fell within 3,974 genes used for differential expression analyses.

We assigned each heterozygous allele as the reference allele, alternate allele, or second alternate allele and matched each allele to its corresponding read depth. This allowed us to identify allele specific expression (ASE) by measuring expression variation between the two haplotypes of each gene distinguished by heterozygous sites. We used a binomal test in R (binom.test) to determine if a heterozygous site showed significantly biased expression of one allele over another (*P* < 0.05; McManus et al. 2010; Mack and Nachman 2016). We measured ASE across 3,974 genes expressed in parental species and hybrids. A gene was considered to show ASE in hybrids if a phased heterozygous allele within that gene showed consistent biased expression in all hybrid samples (17-20 dpf n = 4; 8 dpf n = 3) and did not show ASE in any of the parental samples (n = 12 for both developmental stages).

Gene expression controlled by compensatory variation in parental species is often associated with misexpression in their hybrids (Landry et al. 2005, 2007; Bedford and Hartl 2009; Goncalves et al. 2012). Regulatory elements that have opposite effects on the expression level of a particular gene can compensate for one another to produce an optimal level of gene expression favored by stabilizing selection (Denver et al. 2005; Goncalves et al. 2012). Diverging species can evolve alternate compensatory mechanisms while maintaining similar expression levels (True and Haag 2001). Hybrids of such species would have a mismatched combination of regulatory elements that no longer compensate one another, which is expected to result in biased expression of parental alleles (Wittkopp et al. 2004; Landry et al. 2005). Thus, we identified gene expression controlled by compensatory regulatory variation if a gene 1) did not show differential expression between generalists and molluscivores, 2) showed significant ASE at one or more heterozygous sites in hybrids, and 3) did not show ASE at any site in generalists or molluscivores. Finally, we looked for overlap between genes showing compensatory regulation and genes showing misexpression in hybrids.

## Results

### Differential expression between generalists and molluscivores

We previously found 1,014 genes differentially expressed in whole-larvae tissue between six generalists and six molluscivores collected at 8-10 dpf (McGirr and Martin 2018). Here we compared gene expression in whole-larvae tissue collected at exactly 8 dpf (190-194 hours after fertilization rather than 8-10 dpf) between six generalists and six molluscivores. We found 700 out of 17,723 (3.9%) genes differentially expressed between species (Fig 1C). 235 of the 700 genes were annotated as zebrafish orthologs and used for gene ontology enrichment analyses. Encouragingly, the only significantly overrepresented ontology was skeletal system morphogenesis (GO:0048705) which matched 11 differentially expressed genes (Table S4).

We previously found 120 genes differentially expressed in craniofacial tissue between species at 17-20 dpf (McGirr and Martin 2018). Here we reexamined gene expression in those same individuals using a more conservative threshold for genes to be included in differential expression analyses (where a gene must show >= 10 normalized counts in every sample included in the comparison). As expected, we found fewer genes differentially expressed using this more conservative threshold (81 out of 13,901 (0.6%); Fig 1E). These 81 genes did not show enrichment for any biological process ontologies.

### Hybrid misexpression in whole-larvae tissue

We compared gene expression in whole-larvae tissue collected at 8 dpf from generalist and molluscivore populations (n = 12) with expression in their F_1_ hybrids (n = 3) and found that 370 out of 17,705 genes (2.1%) were misexpressed in hybrids (Fig. 1D). Slightly more genes showed underdominant inheritance (209; 1.2%) than overdominant inheritance (154; 0.89%; Fig. 3A and C). The magnitude of differential expression was higher for genes showing underdominant inheritance than overdominant inheritance (Fig. S4; Wilcoxon rank sum test, *P* = 8.5 × 10^−5^). Of the 370 genes showing misexpression, 138 were annotated as zebrafish orthologs used for gene ontology enrichment analyses. The only significantly overrepresented term was cellular lipid metabolic process (GO:0044255).

**Fig 3.**
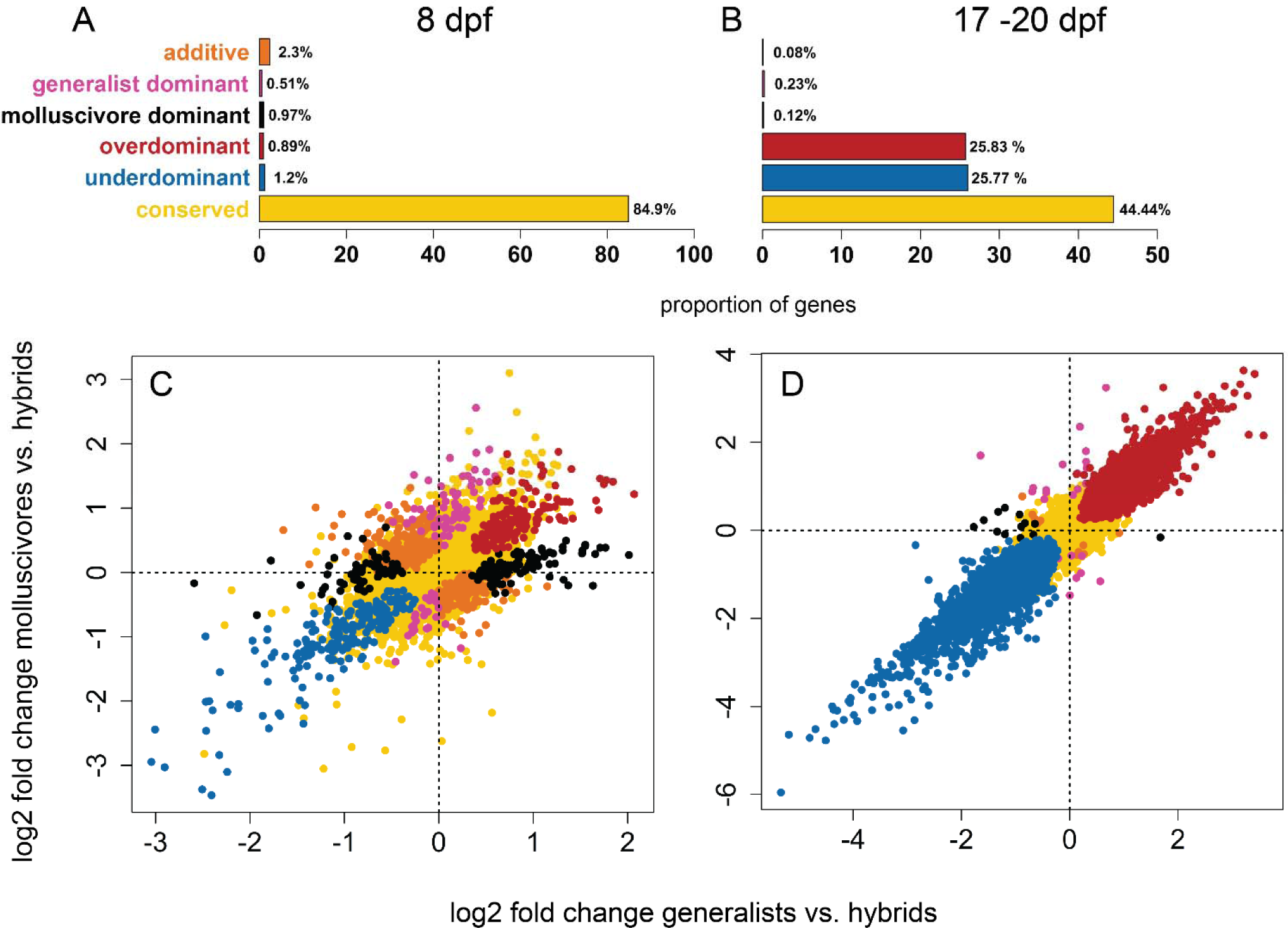
Gene expression inheritance in hybrids. The proportion of A) 17,705 and B) 12,769 genes showing each class of hybrid gene expression in heritance. Log_2_ fold changes in gene expression between molluscivores vs. hybrids on the y-axis and between generalists vs. hybrids on the x-axis for C) 8 dpf whole-larvae samples and D) 17-20 dpf craniofacial samples.

The majority of genes showed conserved levels of expression with no significant difference between hybrids and parental species (84.9%). In line with other hybrid expression studies (Hughes et al. 2006; Rottscheidt and Harr 2007; Davidson and Balakrishnan 2016), most genes that did not show conserved inheritance showed additive inheritance (399; 2.3%). We found some genes showing evidence for dominance, with 89 (0.51%) showing ‘generalist-like’ expression patterns and 168 (0.97%) showing ‘molluscivore-like’ patterns of inheritance (Fig 3A and C).

### Hybrid misexpression in craniofacial tissue

We compared gene expression in craniofacial tissue collected at 17-20 dpf from generalist and molluscivore populations (n = 12) with expression in their F_1_ hybrids (n = 4) and found extensive hybrid misexpression. More than half of genes (6,590 out of 12,769 (51.6%)) were differentially expressed in hybrids compared to parental species expression (Fig 1F). There was an approximately equal number of genes showing overdominant and underdominant expression in hybrids, with 3,299 (25.83%) genes showing higher expression in hybrids relative to parental species and 3,291 (25.77%) showing lower expression in hybrids (Fig 1F, Fig 3B and D). While there was a similar number of genes showing over- and underdominance, the magnitude of differential expression was higher for genes showing underdominance (Fig. S4; Wilcoxon rank sum test, *P* < 2.2 × 10^−16^). Of the 6,590 genes showing misexpression, 2,876 were annotated as zebrafish orthologs used for gene ontology enrichment analyses. Misexpressed genes were enriched for 210 ontologies, including embryonic cranial skeleton morphogenesis (GO:0048701; Table S5 and S6).

### Hybrid misexpression is influenced by library preparation and sequencing conditions

All of the 8 dpf samples were sequenced at the same facility using the same library preparation kit. However, the 17-20 dpf generalist and molluscivore samples were sequenced at a different facility than the 17-20 dpf hybrid samples and used a different library preparation kit. We took two approaches toward understanding how sequencing at different facilities and using different library kits may have affected the proportion of genes misexpressed between hybrids and parental species at 17-20 dpf.

First, we performed another differential expression comparison between whole-larvae hybrids collected at 8 dpf and whole-larvae parental species that we collected for a previous study between 8-10 dpf (McGirr and Martin 2018). The 8 dpf hybrids were sequenced at the same facility with the same library kit as the 17-20 dpf hybrids, while the 8-10 dpf parental species were sequenced at the same facility with the same library kit as the 17-20 dpf parental species. This design mirrored the comparison we used to estimate 17-20 dpf hybrid craniofacial misexpression, but at an earlier developmental stage (Fig. S5). Whereas comparisons between 8 dpf hybrids and parental species sequenced under the same conditions revealed 370 genes (2.1%) misexpressed, comparisons between hybrids and parental species sequenced at different sequencing centers with different library preparation kits suggested that 997 (6%) genes were misexpressed – a 37% increase (Fig. S5). This presents a major caveat to our findings, but does not suggest that all genes showing hybrid misexpression in 17-20 dpf craniofacial tissues are false-positives. Using this estimate of bias to correct for different library preparation methods for our 17-20 dpf samples, we estimate that 19.1% genes were misexpressed in hybrid craniofacial tissue rather than 51.6%.

We also investigated whether a significant amount of between-sample variance in read counts was explained by library preparation method or sequencing date. For each sample we quantified the number of normalized read counts, raw read counts, and raw fastq reads. We also estimated the proportion of duplicate reads out of total mapped reads, the median percent GC content across mapped reads, and the uniformity of coverage across mapped reads (median transcript integrity numbers (TINs)). All of these measures could be influenced by different library preparation methods (Alberti et al. 2014; Van Dijk et al. 2014; KAPA Biosystems 2016). Library preparation method was not associated with differences in the number of normalized read counts or median TINs (Fig. 4 A and B; Welch two sample t-test, *P* > 0.05). When we grouped samples by sequencing date rather than library preparation method, we found that the 17-20 dpf hybrid craniofacial samples (sequenced 6/17) did not show any difference in median GC content, raw read counts, or raw fastq reads compared to samples sequenced on different dates (Fig S6). However, these samples did show lower proportions of duplicate reads, fewer normalized read counts, and lower TINs compared to samples sequenced on all other dates (Fig. 4C-E; ANOVA; *P* < 0.01). TINs quantify the uniformity of coverage across transcripts and are informative as a measure of *in vitro* RNA degradation, which likely suggests that hybrid craniofacial samples experienced more degradation than other samples prior to sequencing. Indeed, lower TIN was significantly correlated with a lower number of normalized counts across samples (Fig. 4F; linear regression; *P* = 2.0 × 10^−5^). Given that hybrid craniofacial samples showed lower TINs and lower normalized counts (Fig. 4C and D), we expected to see a bias toward underexpressed genes in hybrids relative to parental species. Instead, we found approximately the same number of genes overexpressed in hybrids (25.83%) as there were genes underexpressed (25.77%; Fig. 1F and 3B).

**Fig 4.**
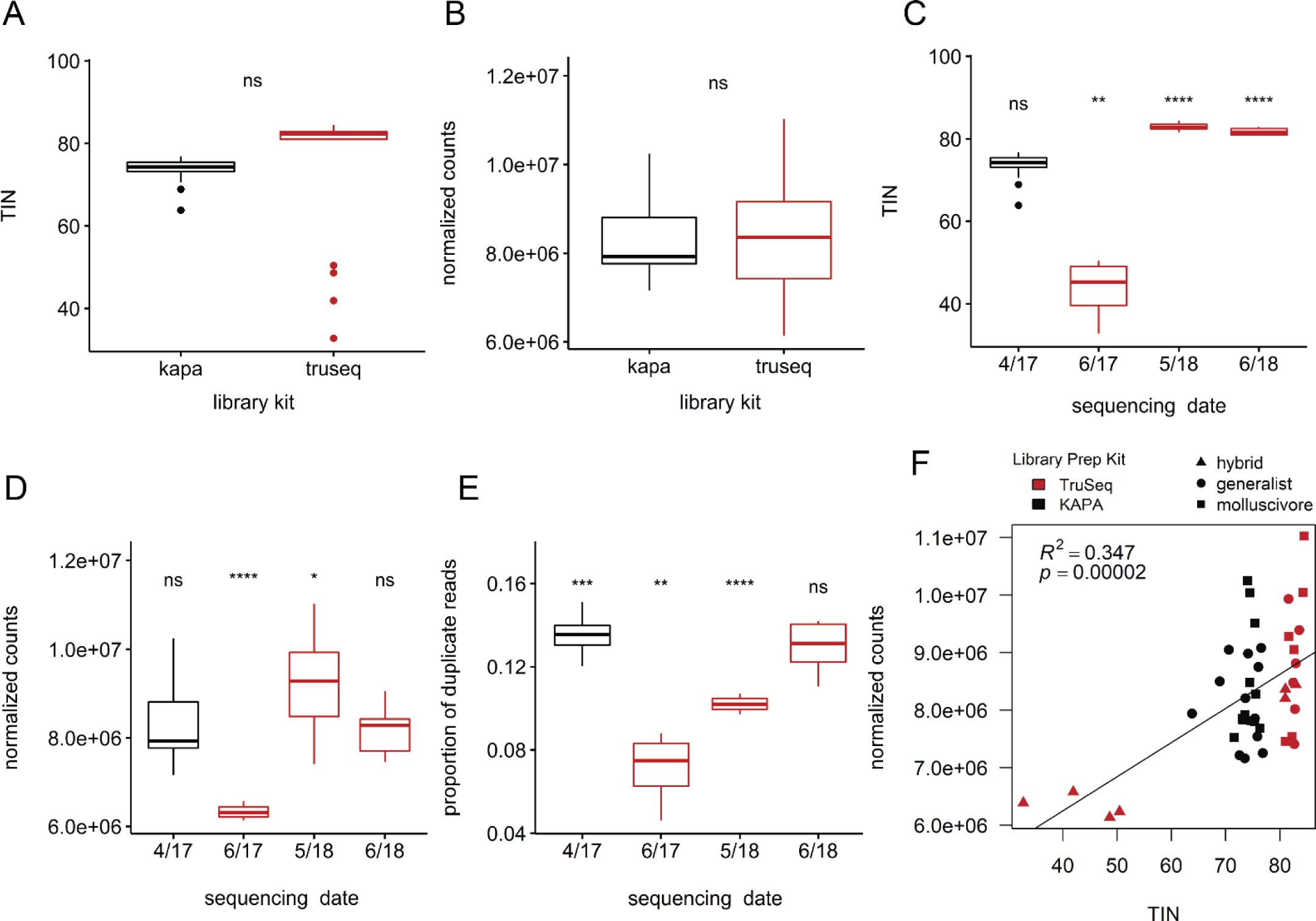
Effects of sequencing facility and library preparation kit. Boxplots show samples grouped by library preparation method (A and B) or by the date they were sequenced (C-E) and whether samples were prepared using Truseq stranded mRNA library kits (red) or KAPA stranded mRNA library kits (black). There was no difference in A) median transcript integrity numbers (TIN) or B) number of normalized counts between groups prepared with different library kits (Welch two sample t-test, *P* > 0.05). 17-20 dpf hybrid craniofacial samples (sequenced 6/17) showed significantly lower C) TIN, D) normalized read counts, and D) proportion of duplicate reads compared to samples sequenced on other dates (ANOVA; *P* < 0.0001 = ****, *** = 0.001, ** = 0.01, * = 0.05). F) Lower TIN was correlated with lower normalized read count.

Overall, we found that our estimate of the proportion of genes misexpressed in 17-20 dpf hybrid craniofacial tissue (51.6%) may be biased due to differences in the number of duplicate reads produced by two different library preparation methods (Fig. 4E). We quantified this bias by measuring hybrid misexpression between samples collected at an earlier developmental stage and found that 19.1% of genes were misexpressed in 17-20 dpf hybrid craniofacial tissues after correcting for library preparation biases (Fig. S5). We found that 17-20 dpf hybrid craniofacial tissues likely experienced more *in vitro* RNA degradation than other samples, but this did not produce a bias toward more genes showing underdominant expression in hybrids (Fig. 3B).

### Compensatory variation underlies misexpression in hybrids

If a gene shows similar gene expression levels between parental species but shows biased allelic expression only in hybrids, it may be regulated by compensatory variation, and such genes are likely to be misexpressed in hybrids (Landry et al. 2005; Goncalves et al. 2012). We identified 15,429 phased heterozygous sites across all 8 dpf and 17-20 dpf individuals with sequencing coverage ≥ 20× that fell within 2,761 (8 dpf) and 1,911 (17-20 dpf) genes used for differential expression analyses. We estimated allele specific expression (ASE) for these genes and paired these data with patterns of differential expression between parental species to identify genes controlled compensatory variation.

We measured ASE across sites within 2,770 genes that showed no difference in expression between generalists and molluscivores at 8 dpf. We found 157 genes (5.4%) that were likely regulated by compensatory mechanisms, which showed ASE only in hybrids and were not differentially expressed between generalists and molluscivores. Of these, nine genes (0.33%) also showed misexpression in hybrids (Fig. 5A and C).

**Fig 5.**
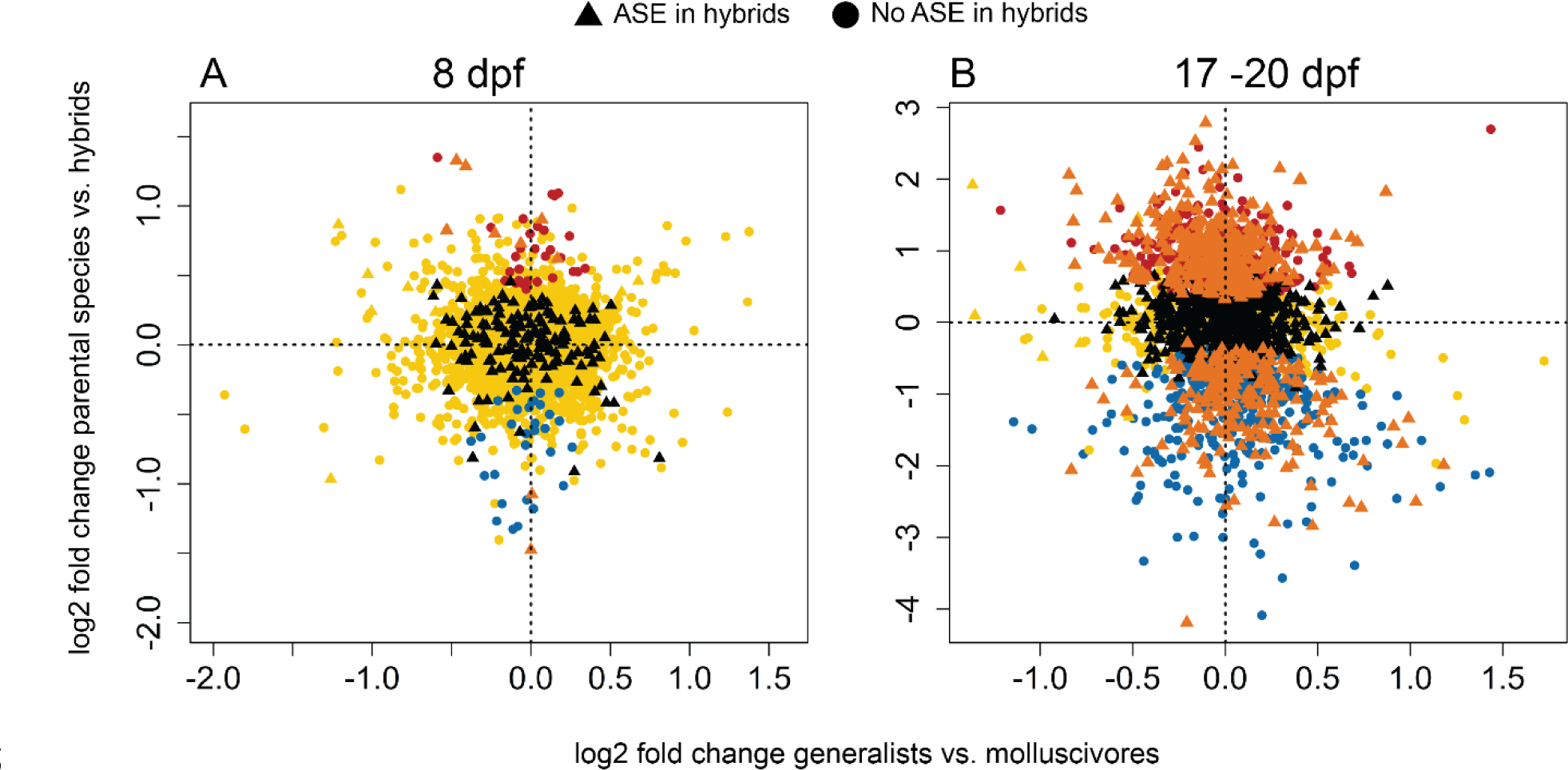
Compensatory regulation underlying expression divergence between generalists and molluscivores. Log_2_ fold changes in gene expression between parental species vs. hybrids on the y-axis and between generalists vs. molluscivores on the x-axis for A) 2,909 genes containing phased heterozygous sites used for allele specific expression (ASE) analyses in 8 dpf whole-larvae samples and B) 2,403 genes containing heterozygous sites in 17-20 dpf craniofacial samples. Triangle points indicate genes showing significant ASE in all hybrids that did not show ASE in generalists or molluscivores. Circle points indicate genes that did not show significant ASE in hybrids or did not show ASE unique to hybrids. Orange = compensatory regulation and hybrid misexpression (genes showing ASE in hybrids, no difference in expression between generalists and molluscivores, and misexpression in hybrids). Black = compensatory regulation (genes showing ASE in hybrids, no difference in expression between generalists and molluscivores). Blue = overdominant (upregulated in hybrids). Red = underdominant (downregulated in hybrids). Yellow = conserved/ambiguous (No difference in expression between parental populations and hybrids).

We also measured ASE across sites within 2,387 genes that showed no difference in expression between generalists and molluscivores at 17-20 dpf. We found 1080 genes (44.81%) that were likely regulated by compensatory mechanisms. In support of this wide-spread compensatory regulation, 581 of these 1080 genes (53.8%) also showed misexpression in hybrids (Fig. 5B and D).

We found many more genes showing ASE in 17-20 dpf hybrid craniofacial tissue than any other samples (Fig. 6A; ANOVA, *P* = 2.81 × 10^−5^). Since misexpression is expected in hybrids when gene expression is controlled by compensatory variation between parental species (Landry et al. 2005; Bedford and Hartl 2009; Goncalves et al. 2012), the high number of genes showing compensatory regulation and high number of genes showing ASE in hybrids supports the validity of extensive misexpression in 17-20 dpf hybrid craniofacial tissue. We likely overestimated the amount of misexpression in this tissue because hybrids were sequenced using a different library preparation kit than parental species (see above). However, ASE was estimated by examining allelic ratios in individual samples and should not suffer from this bias. 17-20 dpf hybrid craniofacial tissue was sequenced at the same facility using the same library preparation kit as all of the 8 dpf samples (Table 1 and S1), yet we only found a high number of genes showing ASE in the 17-20 dpf hybrids (Fig 6A).

**Fig 6.**
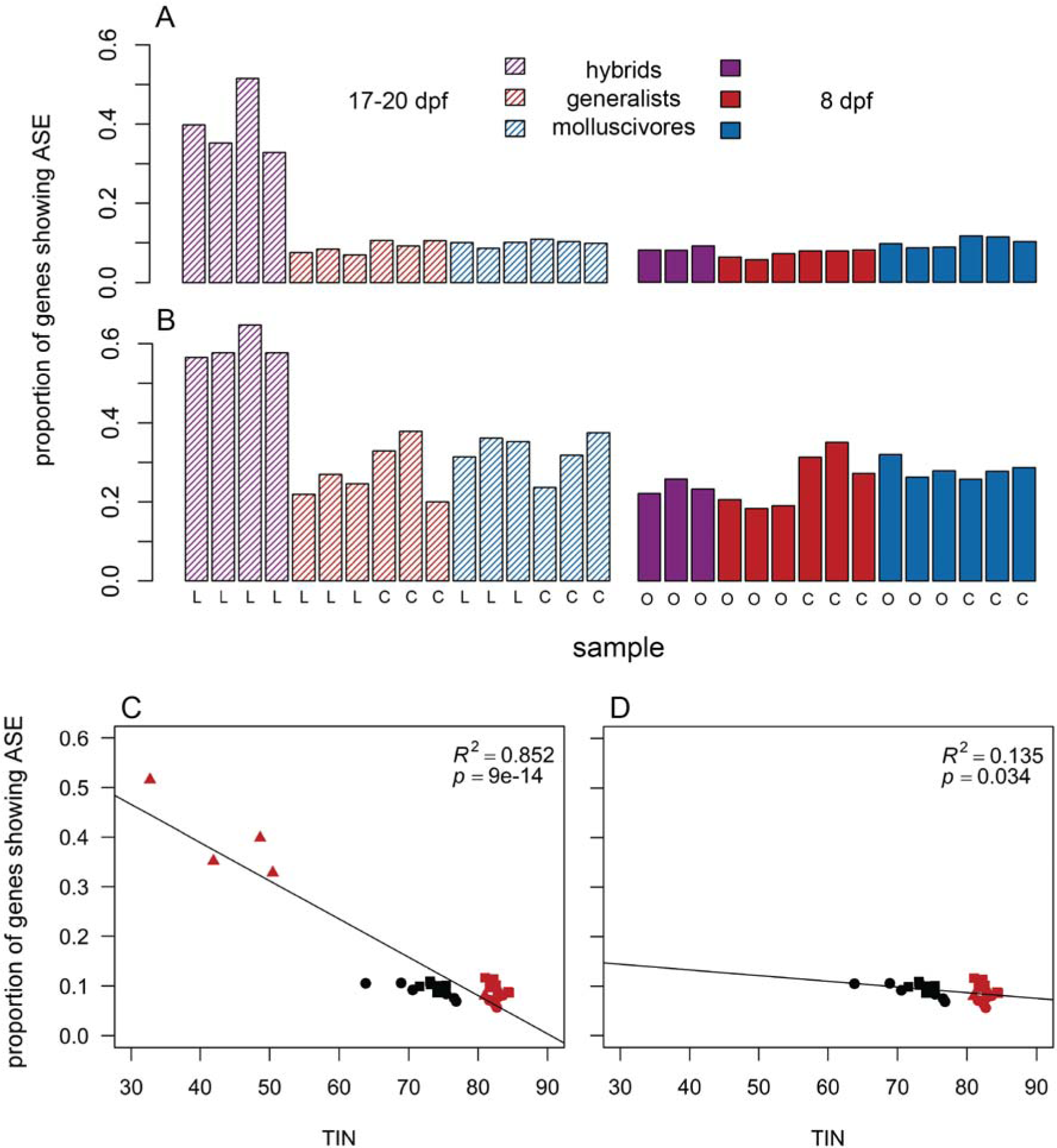
Hybrid craniofacial tissues show high levels of allele specific expression. 17-20 dpf hybrid craniofacial samples (striped purple bars) showed a higher proportion of genes showing significant allele specific expression compared to all other samples using a coverage threshold of A) ≥ 10× reads supporting each heterozygous allele (ANOVA, *P* = 2.81 × 10^−5^) and B) ≥ 100× reads supporting each allele (ANOVA, *P* = 3.85 × 10^−4^). 8 dpf = solid, 17-20 dpf = striped; hybrids = purple, generalists = red, molluscivores = blue; L = Little Lake, C = Crescent Pond, O = Osprey Lake. C) TIN was significantly negatively correlated with ASE (linear regression; *P* = 9.04 × 10^−14^). D) This correlation persisted when 17-20 dpf hybrid craniofacial samples were excluded from the linear model (linear regression; *P* = 0.034). However, the observed proportion of genes showing ASE was much higher in 17-20 dpf hybrid craniofacial samples than predicted by the linear model in D.

We tested whether this pattern might be due to higher rates of *in vitro* degradation in hybrid samples (reflected by low TINs), which could increase variance in the abundance of reads at heterozygous sites and bias ASE estimates. Lower TIN was correlated with higher ASE (Fig. 6D; linear regression; *P* = 9.04 × 10^−14^). This correlation persisted when 17-20 dpf hybrid craniofacial samples were excluded from the model (Fig. 6E; linear regression; *P* = 0.034).

However, the proportion of genes showing ASE was much higher in 17-20 dpf hybrid craniofacial samples than predicted by the latter linear model. Even the lowest TIN for a 17-20 dpf hybrid sample (32.68) predicted a much lower range of genes showing ASE (8.2% −14.1%) compared to the observed range (32.8% - 51.6%). We also estimated ASE again with a higher coverage threshold (>=100 counts supporting each heterozygous allele) to reduce the chances of increased variance affecting binomial tests and still found that hybrid craniofacial samples showed more ASE than other samples (Fig. 6B; ANOVA, *P* = 3.85 × 10^−4^).

## Discussion

Molluscivores show extreme craniofacial divergence relative to their generalist sister species, exhibiting a novel maxillary protrusion and short robust jaws (Fig 1A; Martin and Wainwright 2013a; Hernandez et al. 2018). Given the extreme craniofacial divergence observed between molluscivores and their generalist sister-species, we might expect to find genes expressed in hybrids outside the range of either parent species as a result of discordance between alternatively coadapted genes in networks shaping divergent craniofacial morphologies. However, genetic divergence between generalists and molluscivores is low, with only 79 SNPs fixed between species (genome-wide average F_st_ = 0.08, D_xy_ = 0.00166 (McGirr and Martin 2017; McGirr and Martin 2018)). Despite this low genetic divergence and ongoing gene flow between species, we found misexpression in hybrids at two developmental stages and tissue types. We also measured allele specific expression (ASE) for genes expressed in hybrids and parental species and found evidence for compensatory divergence influencing hybrid misexpression at both developmental stages.

### Hybrid misexpression during juvenile development

While many studies on hybrid misexpression search for regulatory divergence in ‘speciation genes’ associated with sterility and inviability (Malone and Michalak 2008; Renaut et al. 2009; Davidson and Balakrishnan 2016), our results highlight the importance of considering misexpression over multiple early developmental stages and in the context of adaptive morphological traits. We found evidence of misexpression in 8 dpf whole-larvae hybrid tissues (2.1% of genes) and in 17-20 dpf hybrid craniofacial tissues (19.1% of genes after correcting for bias due to library preparation method).

There are several reasons why we might expect to find a higher proportion of genes misexpressed in 17-20 dpf hybrid craniofacial tissues relative to 8 dpf whole-larvae tissues. The molluscivore shows exceptional rates of morphological diversification, particularly in craniofacial traits (Martin and Wainwright 2011). Perhaps 17-20 dpf is a crucial developmental window when gene networks shaping these traits become extensively misregulated in hybrids. It is just after this stage that the relative length of the premaxilla, maxilla, palatine, and lower jaw tend to increase more for generalists than molluscivores (Lencer et al. 2016). It is also possible that regulatory changes are compounded throughout development and have cascading effects, resulting in higher rates of misexpression in later stages. Finally, some of the increased misexpression in hybrid craniofacial tissue can likely be attributed to our sampling design. We found that hybrid craniofacial samples showed lower TINs and lower normalized counts (Fig. 4A and D), suggesting that these samples may have experienced more *in vitro* RNA degradation than other samples (Wang et al. 2016). However, under this scenario, we would expect to see a bias toward lower gene expression in hybrids relative to parental species. Alternatively, we found approximately the same number of genes overexpressed in hybrids (25.83%) as there were genes underexpressed (25.77%), suggesting that many genes were overexpressed in hybrids despite potential RNA degradation.

We found roughly twice the amount of misexpression in hybrid craniofacial tissues compared to a study of misexpression in whole-body tissue that measured gene expression in F_1_ hybrids generated between benthic and limnetic lake whitefish (Renaut et al. 2009). These populations also diverged within the past 10 kya and occupy different habitats within lakes (Renaut et al. 2009; Bernatchez 2004). We also found that genes showing underdominance in hybrids showed a higher magnitude of differential expression compared to those showing overdominance in 8 dpf and 17-20 dpf tissues (Fig. S4), a pattern that has also been observed in whitefish (Renaut and Bernatchez 2011) and a generalist/specialist *Drosophila* species pair (McManus et al. 2010).

### The consequences of hybrid misexpression

It is unclear whether such extensive gene misexpression in hybrid craniofacial tissues might contribute to intrinsic post-zygotic isolation between generalists and molluscivores. F_2_ hybrids exhibiting intermediate and transgressive craniofacial phenotypes showed reduced survival and growth rates in the wild relative to F_2_ hybrids resembling parental species (Martin and Wainwright 2013b; Martin 2016), but short-term experiments measuring F_2_ hybrid survival in the lab did not find any evidence of reduced survival for hybrids with intermediate phenotypes (Martin and Wainwright 2013b). This was interpreted as evidence that complex fitness landscapes measured in field enclosures on San Salvador with multiple peaks corresponding to the generalist and molluscivore phenotypes were due to competition and foraging ability in the wild (i.e. extrinsic reproductive isolation). However, additional analyses of these data suggest that absolute performance of hybrids may also play a role in their survival. The most transgressive hybrid phenotypes exhibited the lowest fitness, contrary to expectations from negative frequency-dependent disruptive selection (Martin 2016). It is still possible that intrinsic and extrinsic incompatibilities interact such that gene misexpression weakens performance more in the wild than in the lab. However, note that F_1_ hybrids used in this study would fall within an intermediate phenotypic position relative to parental trophic morphology whereas field experiments used F_2_ and later generation hybrid intercrosses and backcrosses.

### Hybrid misexpression is controlled by compensatory divergence

When an optimal level of gene expression is favored by stabilizing selection, compensatory variation can accumulate between species and cause misexpression in hybrids (Landry et al. 2005; Bedford and Hartl 2009). We combined results from differential expression analyses with allele specific expression (ASE) results to identify genes controlled by compensatory regulatory divergence between generalists and molluscivores. In 8 dpf whole-larvae tissue, we found 5.4% of genes controlled by compensatory regulation (Fig. 5B). The low amount of genes controlled by compensatory regulation was reflected by the low amount of genes misexpressed in 8 dpf hybrids (2.1%). In 17-20 dpf hybrid craniofacial tissues, we found 44.81% of genes controlled by compensatory regulation (Fig. 5B). The large number of genes controlled by compensatory regulation is consistent with the extensive misexpression observed in hybrid craniofacial tissue, and the majority of genes showing signs of compensatory regulation were also misexpressed in hybrids (53.8%). This independent line of evidence supporting misexpression in 17-20 dpf hybrid craniofacial tissue was not biased by differences in library preparation methods because allele specific expression was estimated by measuring allelic ratios in individual samples. 17-20 dpf hybrid craniofacial tissue was sequenced at the same facility using the same library preparation kit as the 8 dpf samples, yet we only found a high number of genes showing ASE in the 17-20 dpf hybrids (Fig. 6). These results are also in line with studies finding widespread compensatory evolution in other systems with greater divergence times between species (Landry et al. 2005, 2007; Takahasi et al. 2011; Goncalves et al. 2012; Bell et al. 2013; Mack and Nachman 2017; but also see Fraser 2018).

## Conclusion

We found hybrid misexpression in both whole-larvae tissues cranial tissues sampled at early developmental stages. This points to divergent evolution of developmental networks shaping novel traits in the molluscivore. It is unclear whether such misexpression causes intrinsic incompatibilities in hybrids within this recent adaptive radiation. Investigating mechanisms regulating gene expression between generalists and molluscivores that result in hybrid misexpression will shed light on whether the variants shaping novel traits may also contribute to reproductive isolation. Examining misexpression across multiple early developmental stages in the context of developing tissues that give rise to adaptive traits can paint a more complete picture of genetic incompatibilities that distinguish species.

## Data Availability

All transcriptomic raw sequence reads are available as zipped fastq files on the NCBI BioProject database. Accession: PRJNA391309. Title: Craniofacial divergence in Caribbean Pupfishes.

## Acknowledgements

This study was funded by the University of North Carolina at Chapel Hill and the Miller Institute for Basic Research in the Sciences to CHM. We thank Daniel Matute, Emilie Richards, Michelle St. John, and Sara Suzuki for valuable discussions and computational assistance; Emilie Richards for photographs of specimens in Figure 1; The Vincent J. Coates Genomics Sequencing Laboratory at the University of California, Berkeley and the High-Throughout Sequencing Facility at the University of North Carolina, Chapel Hill for performing RNA library prep and Illumina sequencing; the Gerace Research Centre for accommodation; and the Bahamian government BEST Commission for permission to conduct this research.

## Author Contributions

JAM wrote the manuscript, extracted the RNA samples, and conducted all bioinformatic analyses. Both authors contributed to the conception and development of the ideas and revision of the manuscript.

## Competing interests

We declare we have no competing interests.

