## supplemental tables and figures for "Hybrid misexpression in multiple developing tissues within a recent adaptive radiation of *Cyprinodon* pupfishes"

**Table S1.** mRNA sequencing design.

| round | sequencing date | pooled across *n* lanes | library prep kit |
| --- | --- | --- | --- |
| 1 | 4/17 | 1 | KAPA stranded mRNA |
| 2 | 6/17 | 1 | TruSeq stranded mRNA |
| 3 | 5/18 | 1 | TruSeq stranded mRNA |
| 4 | 7/18 | 3 | TruSeq stranded mRNA |

**Table S2.** Read statistics for samples.

| sample | species | stage | sequencing round | library prep kit | raw fastq reads | reads mapped | raw counts | normalized counts |
| --- | --- | --- | --- | --- | --- | --- | --- | --- |
| 1 | hybrid | 17-20dpf | 2 | truseq | 41912228 | 39531780 | 14471030 | 6134242 |
| 2 | hybrid | 17-20dpf | 2 | truseq | 18451756 | 17360214 | 6363816 | 6577234 |
| 3 | hybrid | 17-20dpf | 2 | truseq | 33541230 | 31461875 | 11473464 | 6386857 |
| 4 | hybrid | 17-20dpf | 2 | truseq | 27720328 | 26006609 | 9659367 | 6234856 |
| 5 | generalist | 8dpf | 3 | truseq | 25656702 | 24407630 | 10461159 | 8815459 |
| 6 | generalist | 8dpf | 3 | truseq | 22804982 | 21721330 | 9245634 | 8017330 |
| 7 | generalist | 8dpf | 3 | truseq | 26313696 | 25476498 | 10757268 | 9390970 |
| 8 | molluscivore | 8dpf | 4 | truseq | 38287748 | 36204014 | 15203504 | 7457667 |
| 9 | molluscivore | 8dpf | 4 | truseq | 34288848 | 32578838 | 13434770 | 7536493 |
| 10 | molluscivore | 8dpf | 4 | truseq | 33962768 | 32384092 | 13443902 | 9051218 |
| 11 | generalist | 8-10dpf | 1 | kapa | 23172714 | 17847934 | 6880522 | 7213838 |
| 12 | generalist | 8-10dpf | 1 | kapa | 20575374 | 19452261 | 7933158 | 8206676 |
| 13 | generalist | 8-10dpf | 1 | kapa | 20631366 | 19202893 | 7750909 | 7161844 |
| 14 | generalist | 17-20dpf | 1 | kapa | 20743782 | 18496992 | 6837539 | 8501245 |
| 15 | generalist | 17-20dpf | 1 | kapa | 18728520 | 16361277 | 6040194 | 9051398 |
| 16 | generalist | 17-20dpf | 1 | kapa | 21338994 | 19399691 | 6922698 | 7940092 |
| 17 | molluscivore | 8-10dpf | 1 | kapa | 19100066 | 17430230 | 6789911 | 7921670 |
| 18 | molluscivore | 8-10dpf | 1 | kapa | 19479052 | 17376013 | 6715924 | 7812869 |
| 19 | molluscivore | 8-10dpf | 1 | kapa | 23224142 | 21581058 | 8519810 | 8476166 |
| 20 | molluscivore | 17-20dpf | 1 | kapa | 21012680 | 18765633 | 7182366 | 7853844 |
| 21 | molluscivore | 17-20dpf | 1 | kapa | 20996520 | 19096064 | 7507215 | 7831725 |
| 22 | molluscivore | 17-20dpf | 1 | kapa | 20731964 | 17371497 | 6216825 | 7522140 |
| 23 | generalist | 8-10dpf | 1 | kapa | 26283022 | 23001257 | 8649498 | 8749038 |
| 24 | generalist | 8-10dpf | 1 | kapa | 29483942 | 27652542 | 11273682 | 7540908 |
| 25 | generalist | 8-10dpf | 1 | kapa | 26094366 | 22722751 | 8639044 | 8982363 |
| 26 | generalist | 17-20dpf | 1 | kapa | 23539660 | 21193066 | 8288540 | 9080255 |
| 27 | generalist | 17-20dpf | 1 | kapa | 22989146 | 20041508 | 7630051 | 7855706 |
| 28 | generalist | 17-20dpf | 1 | kapa | 24875424 | 21412781 | 7819750 | 7254103 |
| 29 | molluscivore | 8-10dpf | 1 | kapa | 25828344 | 22266723 | 8306859 | 7798548 |
| 30 | molluscivore | 8-10dpf | 1 | kapa | 25463686 | 22026757 | 7881773 | 7685499 |
| 31 | molluscivore | 8-10dpf | 1 | kapa | 24912808 | 21994615 | 8135992 | 8278350 |
| 32 | molluscivore | 17-20dpf | 1 | kapa | 24703694 | 21871287 | 8360480 | 10038049 |
| 33 | molluscivore | 17-20dpf | 1 | kapa | 21852694 | 18695831 | 6892571 | 10248139 |
| 34 | molluscivore | 17-20dpf | 1 | kapa | 22560226 | 19029742 | 6997928 | 9514273 |
| 35 | generalist | 8dpf | 3 | truseq | 25934770 | 24751899 | 10559980 | 8480276 |
| 36 | generalist | 8dpf | 3 | truseq | 24781078 | 23652972 | 10100401 | 7407245 |
| 37 | generalist | 8dpf | 3 | truseq | 23199342 | 22179263 | 9546582 | 9933070 |
| 38 | molluscivore | 8dpf | 3 | truseq | 25699038 | 24831966 | 10647567 | 9278752 |
| 39 | molluscivore | 8dpf | 3 | truseq | 31456730 | 30017500 | 12699299 | 11025018 |
| 40 | molluscivore | 8dpf | 3 | truseq | 27239292 | 26189352 | 11032426 | 10043336 |
| 41 | hybrid | 8dpf | 4 | truseq | 27989988 | 26774703 | 11506670 | 8202298 |
| 42 | hybrid | 8dpf | 4 | truseq | 26341200 | 25153435 | 10759990 | 8361452 |
| 43 | hybrid | 8dpf | 4 | truseq | 41450864 | 39962577 | 16950855 | 8441280 |

**Table S3.** Quality control statistics for samples.

| sample | species | stage | median TIN | average depth across features | proportion of duplicate reads | median GC content across reads |
| --- | --- | --- | --- | --- | --- | --- |
| 1 | hybrid | 17-20dpf | 48.63 | 169.59 | 6.80 | 46.42 |
| 2 | hybrid | 17-20dpf | 41.90 | 127.99 | 8.15 | 46.50 |
| 3 | hybrid | 17-20dpf | 32.68 | 165.40 | 4.61 | 45.74 |
| 4 | hybrid | 17-20dpf | 50.43 | 138.23 | 8.79 | 45.99 |
| 5 | generalist | 8dpf | 82.94 | 131.20 | 10.19 | 46.82 |
| 6 | generalist | 8dpf | 82.77 | 129.37 | 10.46 | 47.14 |
| 7 | generalist | 8dpf | 83.55 | 125.86 | 10.03 | 47.30 |
| 8 | molluscivore | 8dpf | 81.01 | 139.18 | 14.19 | 46.22 |
| 9 | molluscivore | 8dpf | 82.25 | 128.50 | 14.18 | 46.91 |
| 10 | molluscivore | 8dpf | 82.59 | 125.39 | 13.67 | 48.03 |
| 11 | generalist | 8-10dpf | 72.56 | 157.17 | 13.08 | 46.25 |
| 12 | generalist | 8-10dpf | 73.65 | 145.73 | 13.00 | 45.42 |
| 13 | generalist | 8-10dpf | 73.53 | 140.65 | 13.40 | 46.28 |
| 14 | generalist | 17-20dpf | 68.89 | 144.59 | 13.97 | 45.36 |
| 15 | generalist | 17-20dpf | 70.57 | 134.99 | 14.22 | 46.27 |
| 16 | generalist | 17-20dpf | 63.81 | 155.01 | 13.60 | 44.83 |
| 17 | molluscivore | 8-10dpf | 73.53 | 132.25 | 13.88 | 46.28 |
| 18 | molluscivore | 8-10dpf | 74.69 | 125.74 | 14.05 | 46.78 |
| 19 | molluscivore | 8-10dpf | 74.43 | 142.56 | 12.79 | 45.92 |
| 20 | molluscivore | 17-20dpf | 73.09 | 132.20 | 14.22 | 46.03 |
| 21 | molluscivore | 17-20dpf | 73.17 | 128.74 | 15.12 | 46.81 |
| 22 | molluscivore | 17-20dpf | 71.57 | 138.66 | 13.06 | 47.44 |
| 23 | generalist | 8-10dpf | 76.01 | 140.15 | 12.42 | 46.50 |
| 24 | generalist | 8-10dpf | 75.82 | 154.90 | 12.05 | 45.65 |
| 25 | generalist | 8-10dpf | 74.11 | 146.22 | 12.72 | 46.21 |
| 26 | generalist | 17-20dpf | 76.56 | 129.96 | 14.25 | 45.57 |
| 27 | generalist | 17-20dpf | 75.39 | 136.84 | 13.92 | 45.89 |
| 28 | generalist | 17-20dpf | 76.83 | 127.75 | 13.48 | 45.58 |
| 29 | molluscivore | 8-10dpf | 75.34 | 132.93 | 13.50 | 45.96 |
| 30 | molluscivore | 8-10dpf | 76.29 | 130.14 | 12.95 | 46.38 |
| 31 | molluscivore | 8-10dpf | 75.54 | 131.94 | 13.25 | 46.49 |
| 32 | molluscivore | 17-20dpf | 74.48 | 142.25 | 14.33 | 45.64 |
| 33 | molluscivore | 17-20dpf | 74.08 | 138.28 | 13.73 | 45.90 |
| 34 | molluscivore | 17-20dpf | 75.39 | 129.94 | 13.65 | 46.43 |
| 35 | generalist | 8dpf | 82.43 | 132.68 | 9.94 | 47.27 |
| 36 | generalist | 8dpf | 82.69 | 125.78 | 10.59 | 47.47 |
| 37 | generalist | 8dpf | 81.58 | 136.72 | 9.71 | 46.98 |
| 38 | molluscivore | 8dpf | 81.63 | 135.55 | 9.91 | 47.33 |
| 39 | molluscivore | 8dpf | 84.49 | 125.89 | 10.69 | 47.31 |
| 40 | molluscivore | 8dpf | 84.31 | 118.45 | 10.35 | 47.61 |
| 41 | hybrid | 8dpf | 80.98 | 134.41 | 12.59 | 47.33 |
| 42 | hybrid | 8dpf | 81.02 | 130.78 | 12.12 | 46.98 |
| 43 | hybrid | 8dpf | 82.94 | 142.90 | 11.04 | 47.51 |

**Table S4.** Differentially expressed genes annotated for effects on skeletal system morphogenesis (GO:0048705). This ontology was the only enriched biological process for genes differentially expressed between generalists and molluscivores at 8 dpf (*P* < 0.05; geneontology.org).

| gene symbol | log_2_ fold change | *P* |
| --- | --- | --- |
| *bmp3* | 0.511242 | 0.039704 |
| *chd7* | 0.423654 | 0.047135 |
| *foxe1* | -0.63748 | 0.004896 |
| *gata3* | 0.369094 | 0.043925 |
| *gfpt1* | -0.29543 | 0.039977 |
| *hand2* | 0.639402 | 0.012518 |
| *kat6a* | 0.55044 | 0.000901 |
| *matn1* | 1.144529 | 0.049159 |
| *matn4* | 0.447086 | 0.000203 |
| *mecom* | 0.552098 | 0.023904 |
| *polr1c* | -0.68794 | 0.026325 |

**Table S5.** Misexpressed genes annotated for effects on embryonic cranial skeleton morphogenesis (GO:0048701). This ontology was one of 210 enriched biological processes for 6,590 genes differentially expressed between hybrids and parental species in craniofacial tissue collected at 17-20 dpf (*P* < 0.05; geneontology.org).

| gene symbol | log_2_ fold change | *P* |
| --- | --- | --- |
| *alcam* | 0.610547 | 0.000222 |
| *alx1* | -0.85427 | 0.033755 |
| *bmp3* | 0.583685 | 0.022167 |
| *crispld2* | -0.94461 | 0.000382 |
| *dcaf7* | -0.81576 | 9.25E-09 |
| *egr1* | -1.18846 | 0.039205 |
| *fam20b* | 0.893436 | 3.84E-05 |
| *fgf3* | 0.707254 | 0.026311 |
| *foxe1* | -0.86424 | 0.023926 |
| *fst* | -1.04804 | 0.000342 |
| *gfpt1* | 1.260497 | 1.00E-14 |
| *gnptab* | 0.847555 | 0.000209 |
| *hand2* | -1.56118 | 1.71E-05 |
| *irf6* | -1.14833 | 2.02E-09 |
| *itga8* | -0.48604 | 0.037746 |
| *kat6a* | -1.09119 | 7.80E-05 |
| *kdm6a* | -0.57079 | 0.005865 |
| *kras* | 0.855153 | 2.61E-05 |
| *leo1* | -0.47694 | 0.012396 |
| *mapre2* | 1.453761 | 2.81E-11 |
| *mecom* | -1.08419 | 0.017119 |
| *med12* | -1.45818 | 2.03E-15 |
| *med14* | 0.758638 | 0.000319 |
| *ocrl* | 1.025415 | 4.44E-05 |
| *pak1* | 0.960502 | 0.000228 |
| *pbx4* | 1.852615 | 2.31E-12 |
| *pdgfra* | -0.48287 | 0.048917 |
| *phf8* | 0.493499 | 0.007445 |
| *pitx2* | 0.807376 | 0.014345 |
| *polr1d* | 0.721823 | 0.032707 |
| *rnf2* | -1.12964 | 3.91E-05 |
| *runx3* | 1.718195 | 3.94E-18 |
| *s1pr2* | 0.415986 | 0.024007 |
| *scfd1* | 0.299537 | 0.028434 |
| *sec23a* | -1.43718 | 3.77E-11 |
| *sec24d* | 0.910752 | 0.005705 |
| *sharpin* | -1.43559 | 2.67E-06 |
| *shh* | 1.07417 | 0.004793 |
| *smo* | -0.46599 | 0.039733 |
| *sphk2* | 0.967353 | 7.29E-07 |
| *tfap2a* | 0.81934 | 0.008711 |
| *tshz2* | -0.68911 | 0.000809 |
| *wls* | -1.49603 | 4.15E-07 |
| *wnt4* | 0.983625 | 0.020119 |
| *xylt1* | 0.661423 | 0.009892 |

**Table S6.** Gene ontologies enriched for 6,590 genes misexpressed between hybrids and parental species in craniofacial tissue collected at 17-20 dpf (*P* < 0.05; geneontology.org).

| GO:0002181 | GO:0006417 | GO:0044257 | GO:0150063 | GO:0071840 |
| --- | --- | --- | --- | --- |
| GO:0042255 | GO:0006364 | GO:0033554 | GO:0009056 | GO:0051246 |
| GO:0042273 | GO:0043603 | GO:0034613 | GO:1901575 | GO:0006950 |
| GO:0006402 | GO:0034248 | GO:0070727 | GO:0048562 | GO:0043412 |
| GO:0051236 | GO:0033365 | GO:0016567 | GO:0010605 | GO:0036211 |
| GO:0006412 | GO:0006396 | GO:0030163 | GO:1901137 | GO:0006464 |
| GO:0000956 | GO:0048701 | GO:0046700 | GO:0034622 | GO:0022607 |
| GO:0050658 | GO:0010467 | GO:0008104 | GO:0009790 | GO:0016043 |
| GO:0050657 | GO:1904888 | GO:0046907 | GO:0034654 | GO:0051173 |
| GO:0043043 | GO:0034470 | GO:0090304 | GO:0044267 | GO:0010604 |
| GO:0015931 | GO:0016072 | GO:1901361 | GO:0048598 | GO:0009893 |
| GO:0006401 | GO:0048704 | GO:0071705 | GO:0044260 | GO:0009888 |
| GO:0034976 | GO:0010498 | GO:0044270 | GO:0048568 | GO:0048856 |
| GO:0006403 | GO:0070647 | GO:0034641 | GO:0048880 | GO:0007275 |
| GO:0006366 | GO:0034660 | GO:0006325 | GO:0019438 | GO:0048731 |
| GO:0006605 | GO:0043161 | GO:0044249 | GO:0018130 | GO:0032502 |
| GO:0007034 | GO:0034655 | GO:0051649 | GO:0006082 | GO:0019222 |
| GO:0022618 | GO:0044265 | GO:1901576 | GO:0044237 | GO:0009653 |
| GO:0072594 | GO:0009059 | GO:0001501 | GO:0006807 | GO:0010468 |
| GO:0043604 | GO:0034645 | GO:0009058 | GO:0009887 | GO:0060255 |
| GO:0090150 | GO:0006886 | GO:0006139 | GO:0043170 | GO:0031323 |
| GO:0006518 | GO:0006520 | GO:0043009 | GO:0031324 | GO:0006810 |
| GO:0071826 | GO:0010608 | GO:0006725 | GO:0045935 | GO:0051234 |
| GO:0016197 | GO:0043632 | GO:0009792 | GO:1901135 | GO:0051171 |
| GO:0016579 | GO:0019941 | GO:0046483 | GO:0007423 | GO:0080090 |
| GO:0051169 | GO:0048193 | GO:0033036 | GO:0006508 | GO:0051179 |
| GO:0006913 | GO:1901566 | GO:0018193 | GO:0044238 | GO:0009987 |
| GO:0016570 | GO:0015031 | GO:0051641 | GO:0044281 | GO:0032501 |
| GO:0042254 | GO:0048705 | GO:1901360 | GO:0008152 | GO:0008150 |
| GO:0022613 | GO:0048706 | GO:0051276 | GO:1901362 | GO:0050896 |
| GO:0016569 | GO:0015833 | GO:0006259 | GO:0051172 | GO:0007165 |
| GO:0000398 | GO:0044271 | GO:0002520 | GO:0071704 | GO:0007154 |
| GO:0000377 | GO:0006511 | GO:0044248 | GO:0019538 | GO:0023052 |
| GO:0000375 | GO:0045184 | GO:0010629 | GO:1901564 | GO:0050877 |
| GO:0070646 | GO:0032446 | GO:0048534 | GO:0016192 | GO:0006955 |
| GO:0061919 | GO:0009057 | GO:0030097 | GO:0044085 | GO:0046777 |
| GO:0006914 | GO:0042886 | GO:0019752 | GO:0065003 | GO:0099537 |
| GO:0008380 | GO:0016070 | GO:0071702 | GO:0006996 | GO:0099536 |
| GO:0017038 | GO:0006974 | GO:1901565 | GO:0055114 | GO:0007268 |
| GO:0016071 | GO:0006281 | GO:0009892 | GO:0032268 | GO:0098916 |
| GO:0006397 | GO:0019439 | GO:0043436 | GO:0043933 | GO:0007187 |
| GO:0006457 | GO:0051603 | GO:0001654 | GO:0048513 | GO:0007186 |

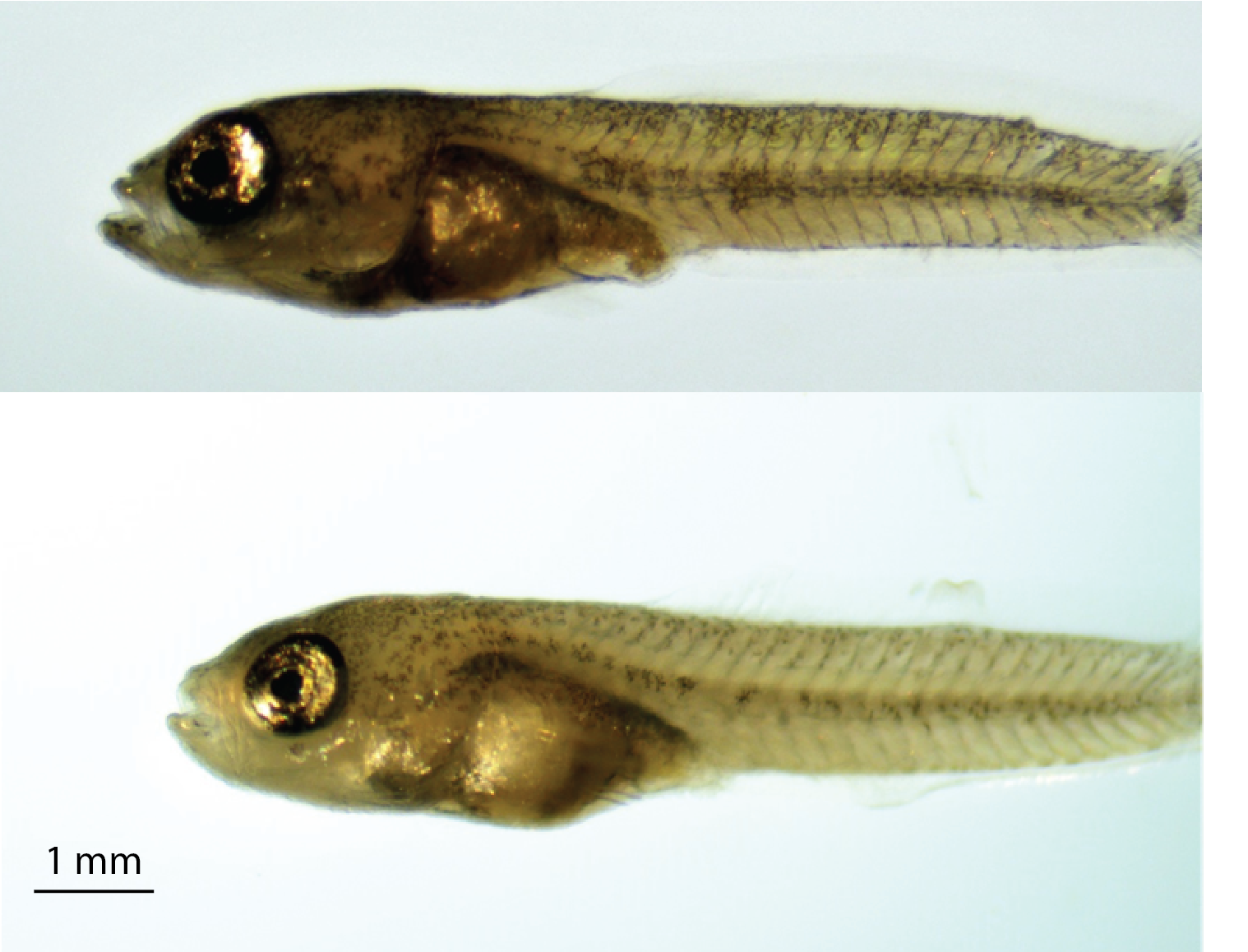

**Fig S1.** 20 day old generalist (top) and molluscivore (bottom).

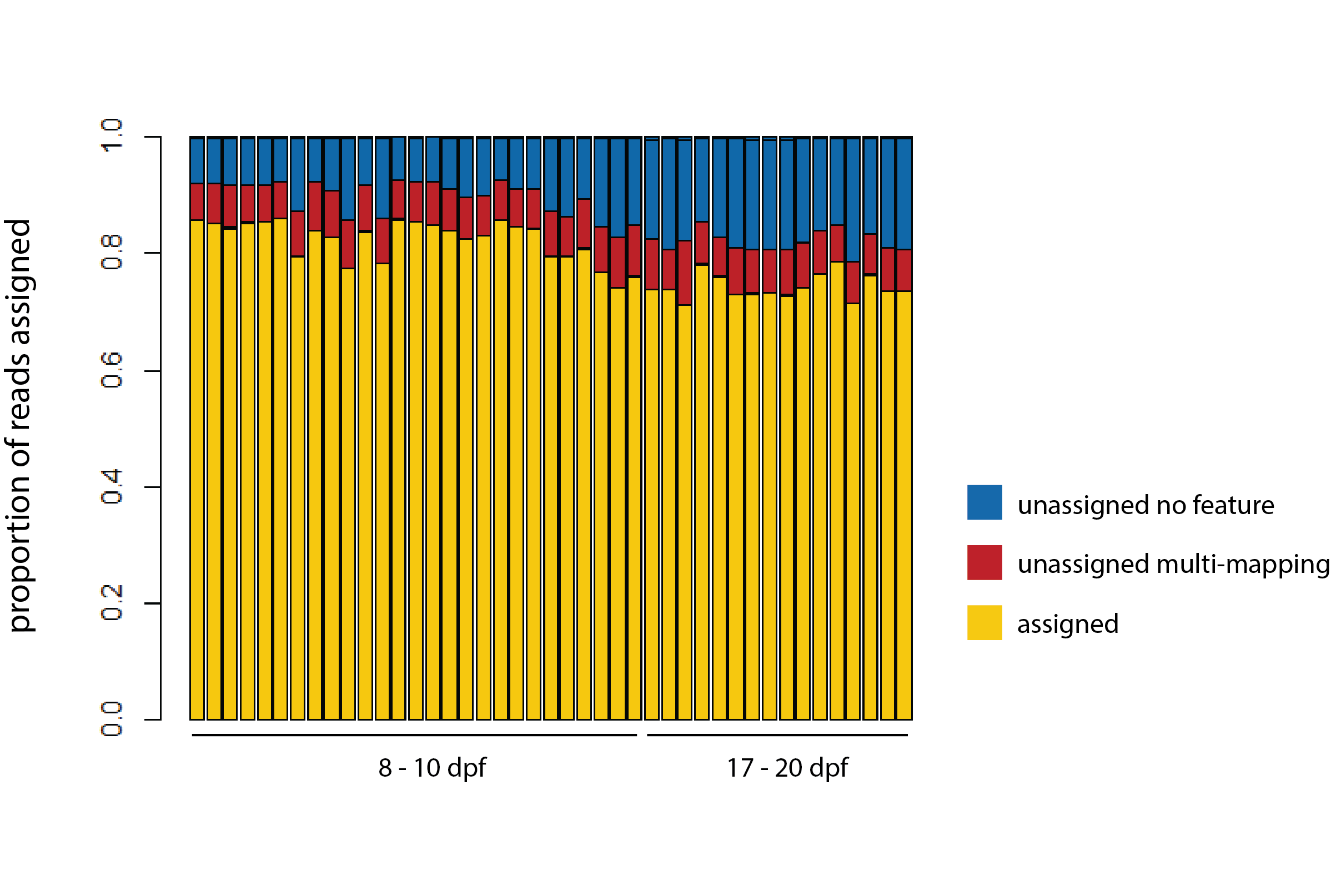

**Figure S2.** Proportion of reads assigned to features (yellow), unassigned due to multi-mapping (red), and unassigned due to no match to annotated features (blue) using STAR aligner.

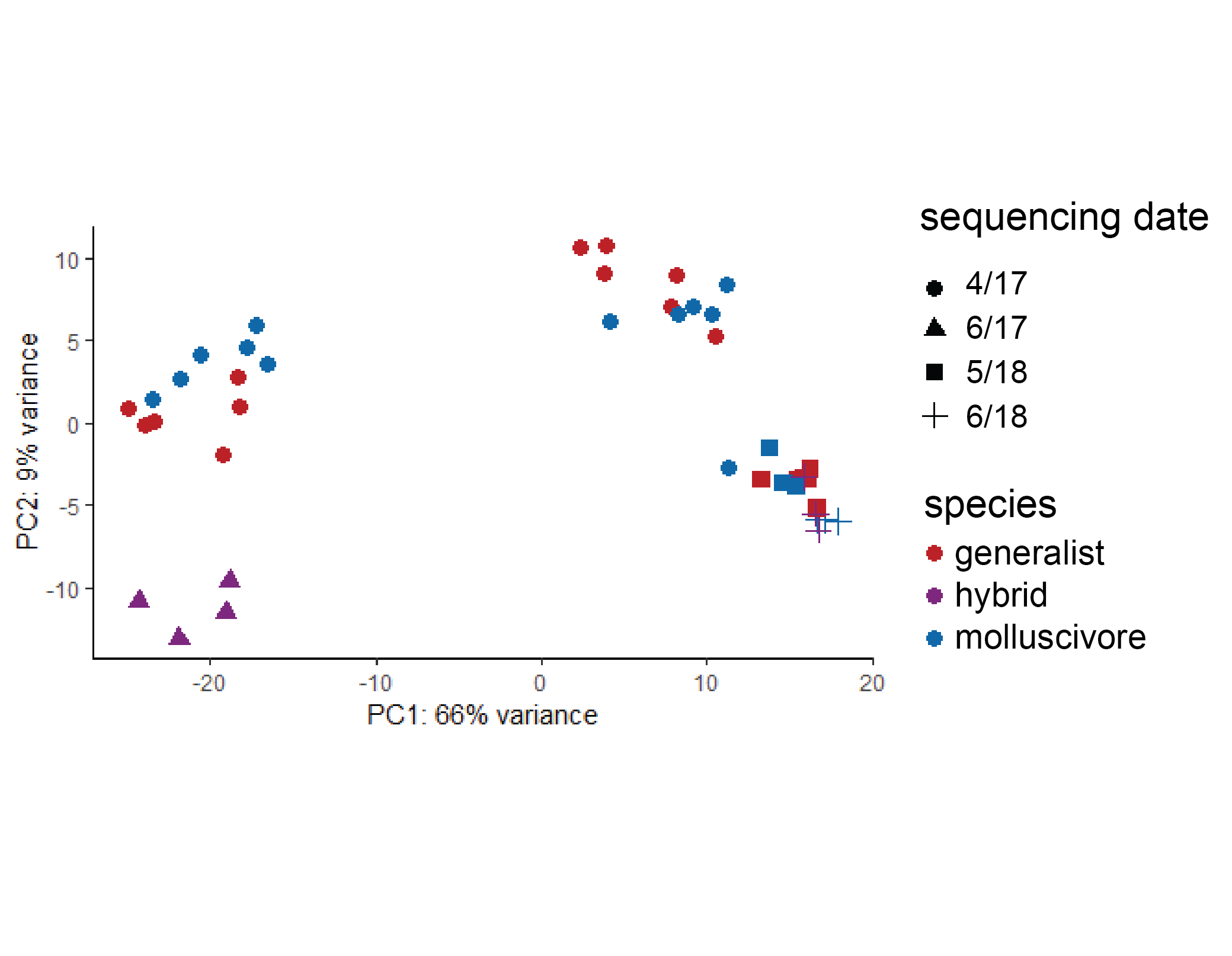
**Figure S3.** The first and second principal component axes accounting for a combined 75% of the total variation between generalist (red), molluscivore (blue), and hybrid (purple) samples across reads mapped to annotated features. Point shape indicates the sequencing date of the sample.

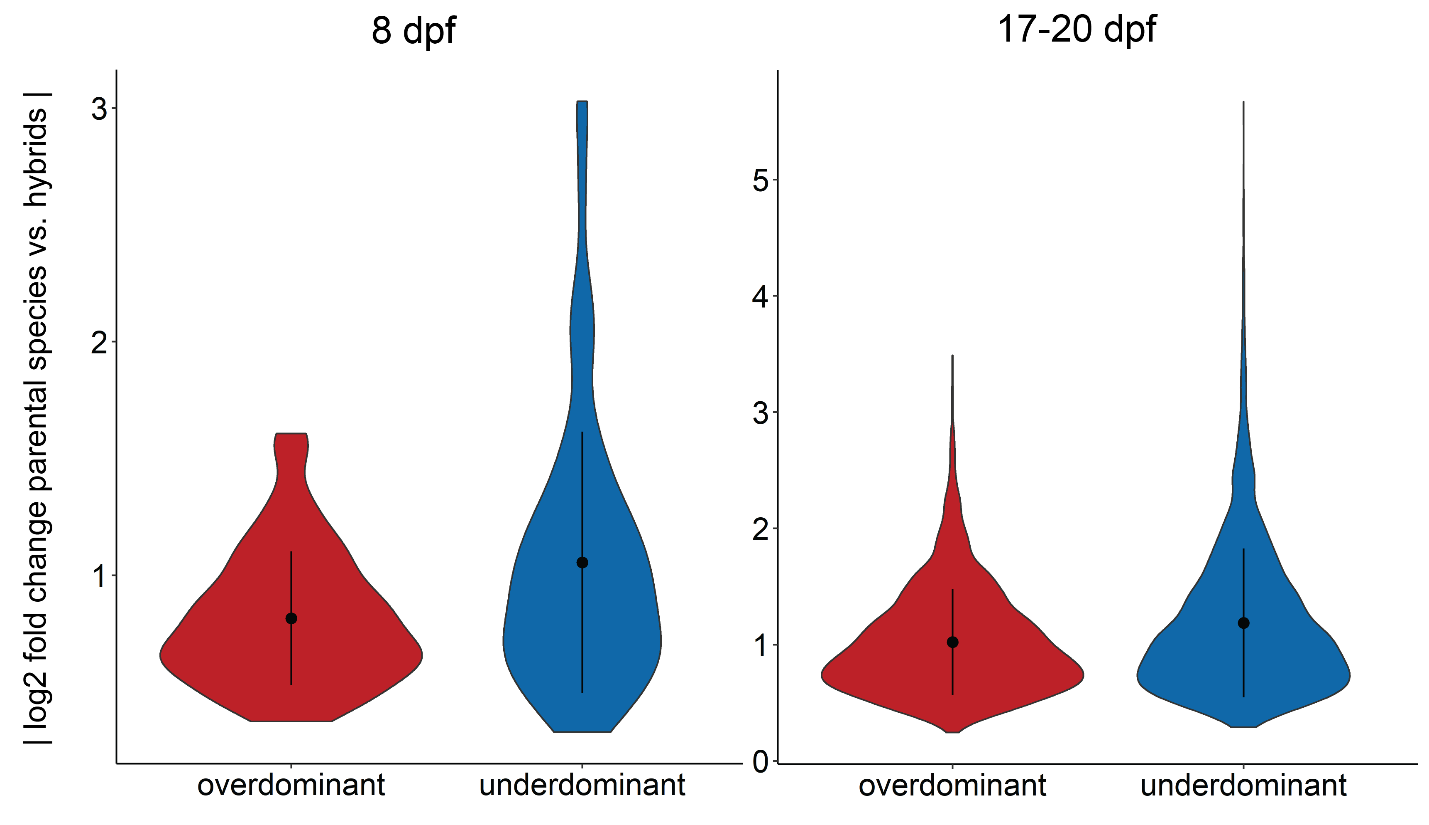

**Fig S4.** Genes showing underdominant expression in hybrids show a higher magnitude of misexpression than genes showing overdominance (Wilcoxon rank sum test; 8 dpf *P* = 8.5 × 10^-5^ 17-20 dpf *P* < 2.2 × 10^-16^).

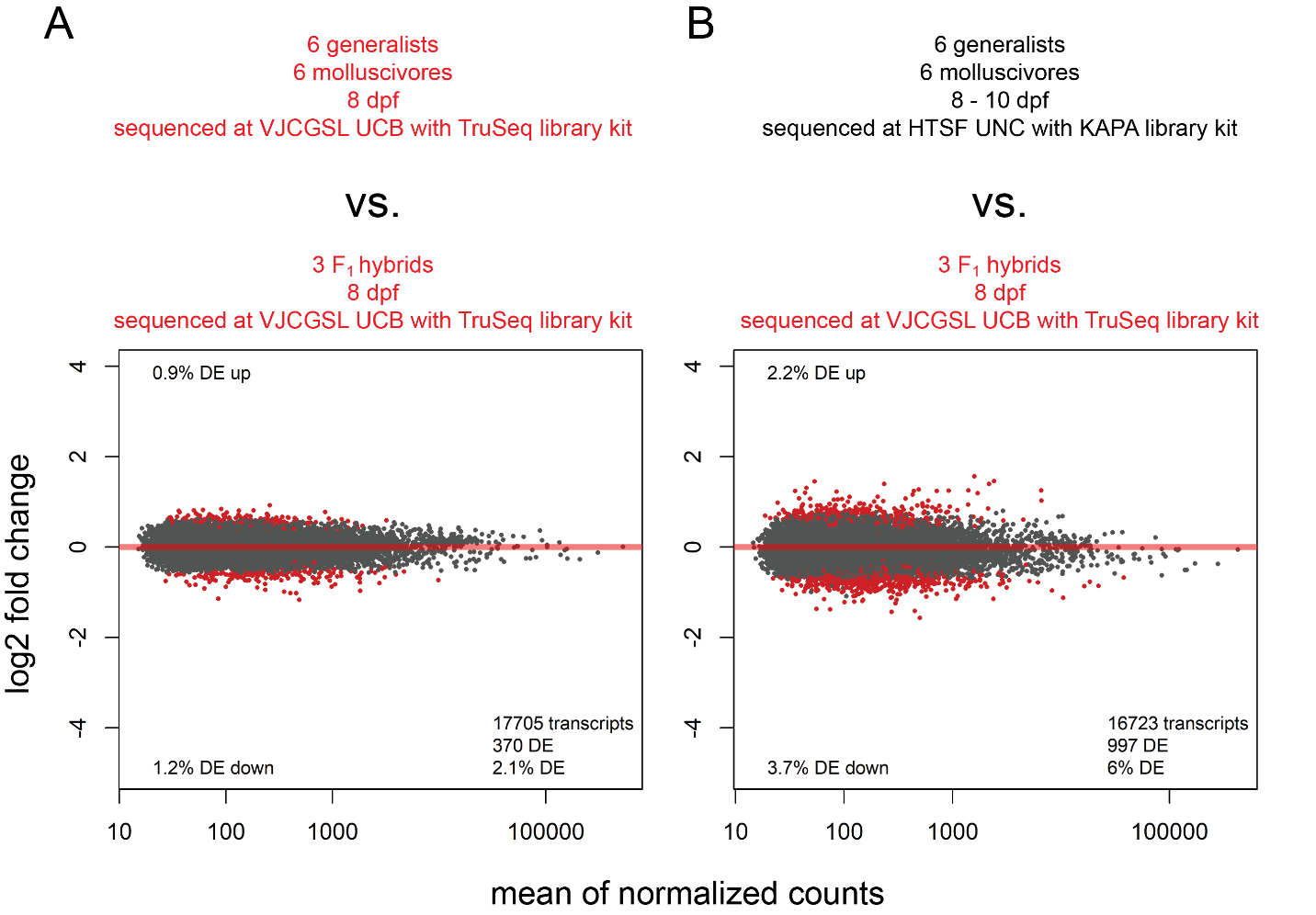

**Fig S5.** **Estimating the effect of sequencing design on the proportion of genes misexpressed in hybrids.** The 8 dpf hybrids were sequenced at the same facility with the same library kit as the 17-20 dpf hybrids, while the 8-10 dpf parental species were sequenced at the same facility with the same library kit as the 17-20 dpf parental species. A) The comparison between 8 dpf parental species and 8 dpf hybrids revealed 370 genes (2.1%) misexpressed. B) The comparison between 8 dpf hybrids and 8-10 dpf parental species revealed 997 (6%) genes misexpressed – a 37% increase. We used this inflated estimate to adjust our estimate of misexpression in 17-20 dpf hybrid craniofacial tissues. Red points indicate genes detected as differentially expressed at 5% false discovery rate with Benjamini-Hochberg multiple testing adjustment. Grey points indicate genes showing no significant difference in expression between groups.

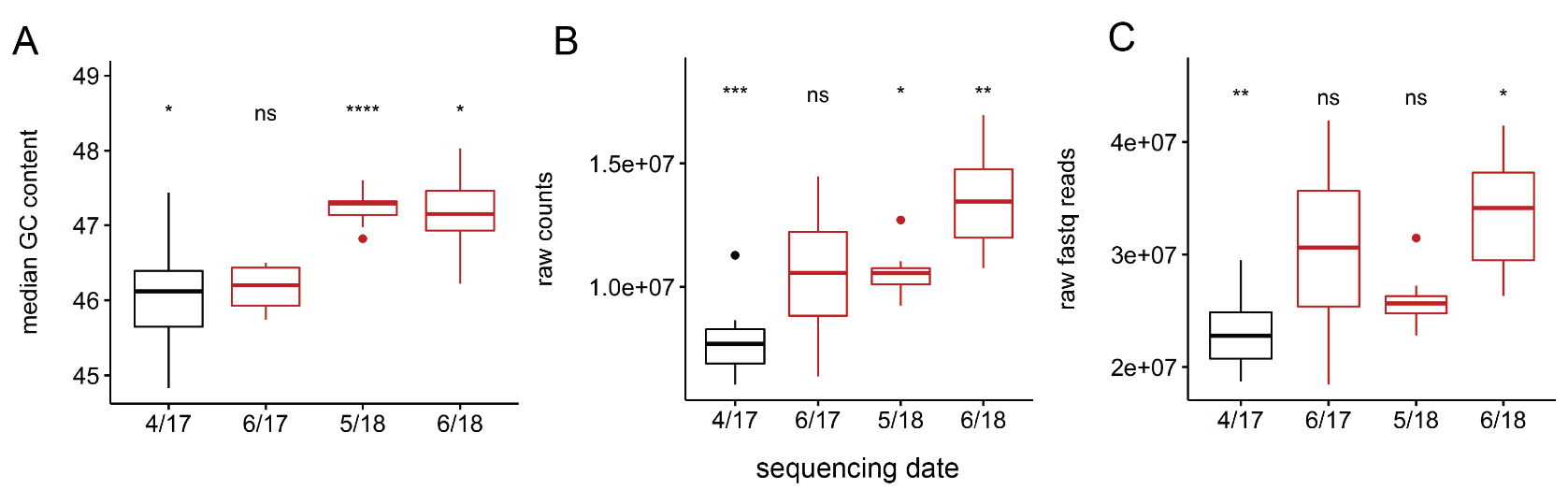

**Fig S6.** We did not find significant differences between 17-20 dpf hybrid craniofacial samples and samples sequenced on other dates for A) median percent GC content across reads, B) number of normalized read counts, or C) number of raw fastq reads. the proportion of duplicate reads for each sample (ANOVA; *P* < 0.0001 = ****, *** = 0.001, ** = 0.01, * = 0.05).

**Coding Supplement**

```
# trim adaptors
trim_galore -q 20 --paired --illumina file1.fastq file2.fastq
# align reads to features
star --runThreadN 4 --genomeDir /genome_dir --readFilesIn file1.fastq file2.fastq --outFileNamePrefix file"
# count reads across features
featureCounts -p -a file.gtf -B -C -G file.fasta -s 2 -T 4 -o files.bam
# haplotype caller
java -Xmx10g -jar GenomeAnalysisTK.jar -T HaplotypeCaller -drf DuplicateRead -R file.fasta -I file.bam  -dontUseSoftClippedBases -stand_call_conf 20.0 -nct 8 -o file.g.vcf
# merge vcfs
vcf-merge file1.g.vcf.gz file2.g.vcf.gz > merged_raw_variants.vcf
# select snps
java -jar GenomeAnalysisTK.jar -T SelectVariants -R file.fasta -V raw_variants.vcf -selectType SNP -o raw_snps.vcf
# filter snps
java -jar GenomeAnalysisTK.jar -T VariantFiltration -R file.fasta -V raw_snps.vcf --filterExpression "QD < 2.0 || FS > 60.0 || MQ < 40.0 || MQRankSum < -12.5 || ReadPosRankSum < -8.0" --filterName "standard_snp_filter" -o filtered_snps.vcf
# filter minor allele frequency 5% and max missing genotype 90%
vcftools --vcf filtered_snps.vcf --maf 0.05 --max-missing 0.9 --recode --out maf_0.5_maxmiss_0.9_snps"
# phase
java -jar GenomeAnalysisTK.jar -T ReadBackedPhasing -R file.fasta -I file.bam --variant snps.vcf -o phased.vcf --phaseQualityThresh 20.0
# count allele reads
java -jar GenomeAnalysisTK.jar -R file.fasta -T ASEReadCounter -U ALLOW_SEQ_DICT_INCOMPATIBILITY -o allele_counts.csv -I  file.bam -sites phased.vcf
# output snp genotypes
java -jar GenomeAnalysisTK.jar -R file.fasta -T VariantsToTable -V phased.vcf -F CHROM -F POS -GF GT -GF HP -o snp_table.txt
# Basic DESeq2 design
  dds <- DESeqDataSetFromMatrix(countData = cts,
                                colData = colData,
                                design= ~species)
  dds <- estimateSizeFactors(dds)
  idx <- rowSums(counts(dds, normalized=TRUE) >= 10 ) >= (nrow(colData))
  dds <- dds[idx,]
  dds <- DESeq(dds)
```
